# A semi-field system for quantifying *Anopheles gambiae* attraction to human scent

**DOI:** 10.1101/2022.12.25.521702

**Authors:** Diego Giraldo, Stephanie Rankin-Turner, Abel Corver, Genevieve M. Tauxe, Anne L. Gao, Dorian M. Jackson, Limonty Simubali, Christopher Book, Jennifer C. Stevenson, Philip E. Thuma, Andrew Gordus, Monicah M. Mburu, Edgar Simulundu, Conor J. McMeniman

**Author notes:** These authors contributed equally (Diego Giraldo, Stephanie Rankin-Turner).

## Abstract

Variability in the chemical composition of human scent has the potential to modulate mosquito attraction to certain humans. We have engineered a large-scale, semi-field system in Zambia for quantifying mosquito olfactory preferences towards whole body odor sourced from different humans under naturalistic conditions. In a flight cage arena with infrared tracking, we document that the African malaria mosquito *Anopheles gambiae* hierarchically prefers to land on heated targets mimicking human skin temperature when they are baited with carbon dioxide (CO_2_) over background air, human body odor over CO_2_, and the scent of one individual over another. In a six-choice assay configuration, we further identify humans at both ends of the attractiveness spectrum whose scent is differentially attractive to *An. gambiae* relative to other individuals. We demonstrate integrative use of this multi-choice olfactory assay with whole body volatilomics, establishing a powerful method for discovery of human odorants modulating heterogeneity in biting risk at enhanced throughput.

## Introduction

The African malaria mosquito *Anopheles gambiae* is a prolific vector of malaria throughout sub-Saharan Africa, a disease that is responsible for over 600,000 deaths worldwide annually ^1^. Malaria transmission occurs between humans via the infectious bite of female anopheline mosquitoes. Several studies have shown that there can be substantial variability in mosquito biting frequency towards humans in disease endemic regions ^2^, and that some people are more attractive to malaria vectors relative to others ^3–8^. Heterogeneity in mosquito biting preferences can lead to variation in infection risk and intensify the basic reproductive number of malaria, which can have important consequences in malaria epidemiology and control ^2, 9^.

Different factors have been associated with variability in human attractiveness to mosquitoes. Amongst these are pregnancy ^8, 10, 11^, skin microbiota ^12–14^, diet (Paskewitz et al., 2018), alcohol consumption ^15, 16^, stage-specific malaria parasite infection ^18–20^, and human genetics ^21, 22^. These factors are all believed to influence the chemical composition of human body odor and associated behavioral responses of mosquitoes towards certain humans ^14, 19^.

To detect humans from a distance, female mosquitoes track a complex blend of airborne chemicals emitted by the human body in skin odor and breath. A number of human-derived chemicals have been identified that when combined together elicit attractive behavioral responses in highly anthropophilic mosquito species including *An. gambiae* and the yellow fever mosquito *Aedes aegypti*. Such compounds include carbon dioxide (CO_2_) ^23, 24^, lactic acid ^25, 26^, ammonia ^27, 28^, and numerous carboxylic acids ^29^. Many of these have been incorporated into synthetic chemical lures that aim to mimic human scent to improve the efficiency of traps for mosquito surveillance and control, such as the BG lure ^30^ consisting of ammonia, hexanoic acid and lactic acid; MB5 lure ^31^ comprising ammonia, lactic acid, tetradecanoic acid, 3-methyl-1-butanol and 1-butylamine; and the Ifakara blend ^32^ made from ammonia, lactic acid, propionic acid, butanoic acid, pentanoic acid, methylbutanoic acid, heptanoic acid, octanoic acid and tetradecanoic acid; all of which are presented concurrently with CO_2_ as a behavioral synergist.

Although many other compounds have been identified as having potential neurophysiological or behavioral activity across various mosquito species ^33^, the attractiveness of the vast majority of volatile organic compounds (VOCs) emitted by the human body remains unknown. Furthermore, the ratio-specific constitution of chemical blends emitted by the human body that modulate preference for different humans, and the biological factors resulting in variation in human odor chemistry are still poorly understood. Although it is well known that the lower extremities including the legs and feet of seated humans and their associated odors are highly attractive to *An. gambiae*, this mosquito species distributes randomly along the lower margins of the body adjacent to the ground when humans lay down – including the head, trunk, arms and legs ^34, 35^. This suggests that convection currents play a primary role in dictating preferred biting sites for *An. gambiae*, and this mosquito species may more broadly track odorants from the entire human body in combination with convection currents to evoke context-specific landing on humans.

Existing behavioral assays to quantify mosquito host preference commonly rely on the use of two or three choice assays with entry or exit traps, or employ decoy traps or electrocution traps whereby mosquitoes are collected onto adhesive surfaces or electrocution grids for quantification ^5, 13, 18, 20, 37, 42–44^. However, whilst highly useful for field-based surveillance activities or small-scale semi-field assays to measure mosquito host preference, these are inherently by design end-point assays, and currently do not facilitate temporal quantification of mosquito landing preferences during host-choice assays. Furthermore, in existing assay formats, mosquito behavioral attraction is often quantified towards odor sourced from specific body parts like arms and feet, with scent being commonly captured onto socks, nylon mesh or glass beads for use in mosquito behavioral testing and downstream chemical analysis ^13, 19, 37, 45, 46^. This results in mosquitoes being exposed only to a subset of compounds present in human scent, and not the full complex blend of airborne compounds emitted by the whole body, potentially limiting the size of chemical space surveyed for discovery of human odorants modulating mosquito behavior.

*Anopheles gambiae* classically exhibits peak periods of host-seeking activity during the late evening and early mornings in the hours flanking midnight, when humans are typically sleeping ^47^. During this time frame, humans are usually indoors, where human whole body odor inclusive of carbon dioxide (CO_2_) and other volatile organic compounds found in skin odor and breath ^48^, dissipates from the human body into the air. Human body odor rises on warm convective currents emanating from the skin to generate odor plumes that typically dissipate through the housing eaves ^49^ or walls to be carried downwind to provide an alluring olfactory stimulus for mosquitoes in the vicinity that indicates human presence ^50–52^. Anthropophilic mosquitoes including *An. gambiae* and *Aedes aegypti* use multiple sensory cues including CO_2_, other constituent body odorants, heat, moisture and vision to detect human hosts ^23, 46, 53, 54^, and these cues often interact synergistically to increase attraction and landing on a surface ^23, 46, 55, 56^.

Towards identifying chemical features that underlie inter-individual differences in human attractiveness to *An. gambiae*, we have developed a large-scale, semi-field system in Zambia for multichoice assays of mosquito olfactory preference towards whole body odor sourced from different humans. Using infrared video tracking, we demonstrate that *An. gambiae* hierarchically prefers to land on warmed targets when they are baited with CO_2_ over background air, human body odor over CO_2_, and the scent of one individual over another. Using a six-choice assay configuration, we further use this system to screen for individuals whose scent is differentially attractive to *An. gambiae*, and demonstrate application of whole body volatilomics to identify human odorants with the potential to modulate attractiveness to this major disease vector. These data indicate that this semi-field system has potential to form the basis of an innovative resource for large-scale screens to identify humans that are highly attractive or unattractive to *An. gambiae* and other malaria vectors, to define the chemosensory basis of malaria transmission.

## Results

### Development of the Odor-Guided Thermotaxis Assay to quantify *Anopheles gambiae* landing behavior under low-light conditions

To generate an optimized assay configuration to mimic the process of landing on human skin, with the potential to be scaled for use in mosquito multi-choice host preference assays, we first engineered a novel behavioral apparatus customized for scoring *An. gambiae* landing behavior in the presence of heat and olfactory stimuli under low-light conditions. The Odor-Guided Thermotaxis Assay (OGTA) uses infrared videography to quantify landings of female *An. gambiae* on a black aluminum disc heated to mimic human skin temperature (35°C), that can be baited with any target olfactory stimulus using controlled airflow (Fig. 1). We designed all OGTA components to be powered using 12-volt batteries to facilitate high-content recordings of mosquito landing behavior in “off-the-grid” settings typically encountered under semi-field and field environments situated at remote localities in Africa.

**Figure 1.**
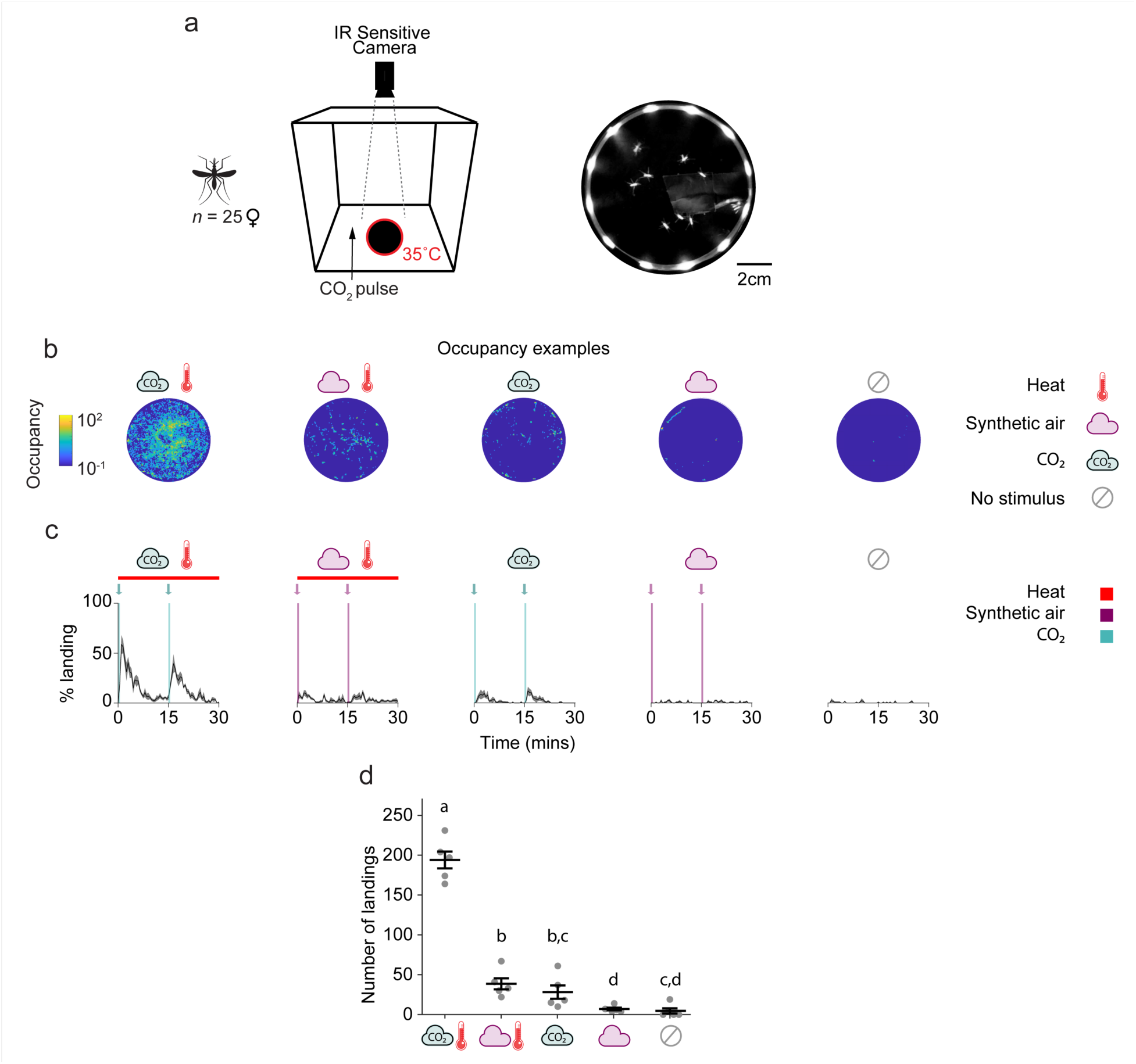
Odor-guided thermotaxis assay (OGTA) for measuring *Anopheles gambiae* landing behavior at night. (a) Schematic of the OGTA for use in laboratory conditions. Twenty-five *An. gambiae* females were introduced into a cage containing a landing platform heated to 35°C surrounded by infrared LEDs. CO_2_ was puffed into the cage and mosquito landings on the platform were recorded using an infrared-sensitive camera throughout a 30 min assay period. (b) Heatmaps of platform occupancy from a representative experiment for each stimulus combination. (c) Percentage of mosquitoes landing on the platform throughout the experiment for each stimulus combination. The arrows indicate the timepoints of stimulus onset. Bars represent the length of stimulus duration (green = CO_2_, purple = synthetic air, two 1-min pulses of each olfactory stimulus). Mean ± SEM plotted. (d) Total number of landings on the platform for each stimulus combination. Mean ± SEM plotted. n = 5/stimulus combination. Fisher’s exact permutation tests with Benjamini-Hochberg correction (a-b: p < 0.05, a-c: p < 0.05, a-d: p < 0.05, b-c: p < 0.05, b-d: p < 0.05, c-d: p < 0.05).

To validate the utility of the OGTA to score odor-guided thermotaxis from *An. gambiae*, we first performed behavioral trials examining multimodal synergisms between the host-related cues CO_2_ and heat in the laboratory, whilst simulating night-time light conditions. Using infrared videography, we determined that female mosquitoes were readily visualized during landings on the infrared illuminated OGTA platform (Fig. 1a). Strikingly, when CO_2_ and heat were co-presented as stimuli in these assays, high levels of landings were observed (Fig. 1b-c and Fig. S1).

In particular, each pulse of CO_2_ triggered significant levels of landings on the heated platform that returned to baseline levels within 15min (Fig. 1b-c and Fig. S1). In contrast, trials with a synthetic air stimulus with heat triggered very few landings. Similarly, CO_2_ pulses without heat yielded a similarly low number of landings. Control assays with synthetic air without heat, or no stimuli at all, elicited even lower levels of landings. We conclude that the OGTA is a powerful method to quantify *An. gambiae* landing behavior under low-light conditions, and that CO_2_ and heat potently synergize together to evoke landing behavior in this primary malaria vector (Fig. 1d).

### A semi-field system for multi-choice assays of mosquito olfactory preference

To develop an expansive arena for multi-choice assays of *An. gambiae* olfactory preference with arrays of OGTAs in competition, we next engineered and constructed a large-scale, semi-field system at Macha, Choma District, Zambia (Fig. 2). This system consists of a central screened flight cage (20m x 20m x 2.5m) with a volume of 1000m^3^ for contained assays of *An. gambiae* behavior that is flanked at its perimeter by up to eight one-person tents (Fig. 2a-b). Each tent is connected to the central cage by screened aluminum ducting with a low-speed fan to directionally pipe target olfactory cues placed inside them, including whole body odor from sleeping humans or CO_2_, into the flight cage arena and directly onto concordant OGTAs (Fig. 2c and Fig. S3) for exposure-free assays of mosquito olfactory preference. For use in the semi-field system, we modified the OGTA design to place the infrared-illuminated landing platform heated to 35°C and IR-sensitive camera on top of a standardized black container, with the screened ducting from each tent venting odor directly over this target (Fig. 2d).

**Figure 2.**
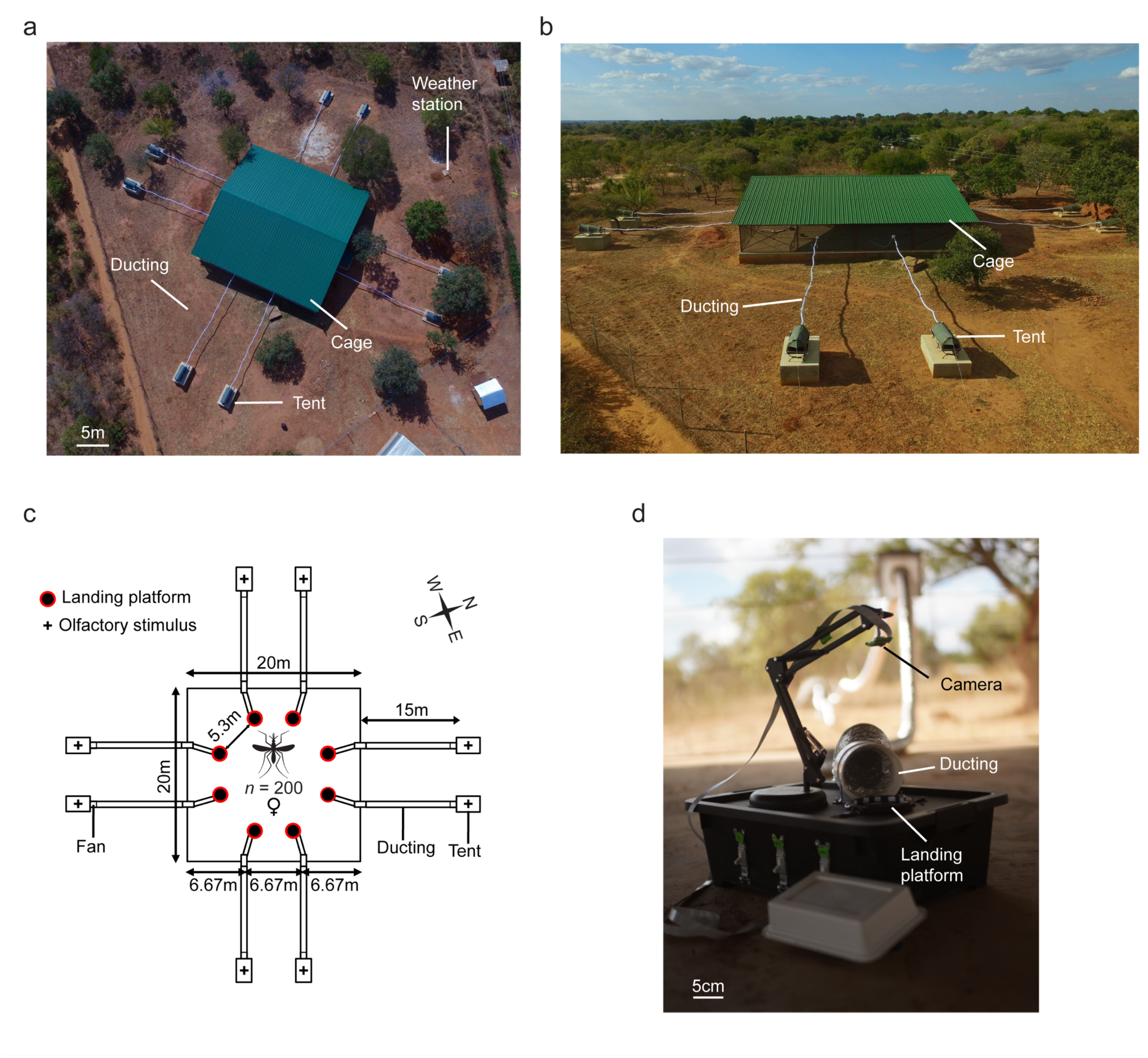
Semi-field system for multi-choice assays of mosquito olfactory behavior. (a) Aerial and (b) side views of the screened mosquito flight cage arena (volume: ∼1000m^3^) that is surrounded by 8 one-person tents. The facility has a roof for protection of the cage and assay components from inclement weather (green). Tents (2 per side) are positioned 15m away from the cage and are connected to the central cage by ducting, to deliver olfactory stimuli via a ducting fan assembly to the flight cage arena. A weather station outside of the cage collects weather data. (c) Schematic of the multi-choice olfactory preference assay. (d) The ducting from each tent vents the target olfactory stimulus over an infrared-illuminated landing platform heated to 35°C mimicking human skin temperature, and mosquito landings are recorded throughout the duration of each assay using an infrared-sensitive camera.

This semi-field system provides flexibility to perform anywhere from one-choice to eight-choice trials, by simply modifying the number of tents and OGTAs used in experiments. In contrast to typical two-choice laboratory wind tunnels with volumes of ∼0.5m^3^ (example dimensions: 2m x 0.5m x 0.5m), our semi-field system provides a flight cage arena that is ∼ 2000-fold larger to assess *An. gambiae* olfactory preferences under naturalistic conditions. Conveniently, separate insectaries located adjacent to the semi-field system facilitated use of laboratory-reared Kisumu strain *An. gambiae* of known age and physiological status for behavioral experimentation. A weather station (Fig. 2a) was also installed adjacent to the flight cage to measure environmental parameters including wind speed and directionality, temperature and humidity; and retrospectively monitor their effects on *An. gambiae* host-seeking behavior.

### *Anopheles gambiae* prefers heated targets baited with CO*_2_* relative to background air

To initially validate the utility of this semi-field system to quantify mosquito olfactory preferences, we first tested choices of female *An. gambiae* to land on one heated OGTA platform baited with a CO_2_ stimulus that was 400ppm above background atmospheric concentrations, relative to seven heated platforms that were baited with only background air (Fig. 3, Fig. S2 and Fig. S4). For these trials, the eight OGTAs were arrayed in an octagonal configuration in the center of the flight cage arena (Fig. 2c and Fig. S3), with 6 replicate nightly assays occurring from 22:00-4:00 hours (Fig. 3a and Fig. S2a), to occur during peak periods of *An. gambiae* circadian activity. Consistent with our previous laboratory assays (Fig. 1), we observed a strong synergy between CO_2_ and heat under semi-field conditions, with these cues eliciting high levels of landings only on the platform where these stimuli were co-presented (Fig 3b-c, Fig. S2b and Fig. S4). In contrast, very few landings were observed on the other seven heated platforms baited only with background air.

**Figure 3.**
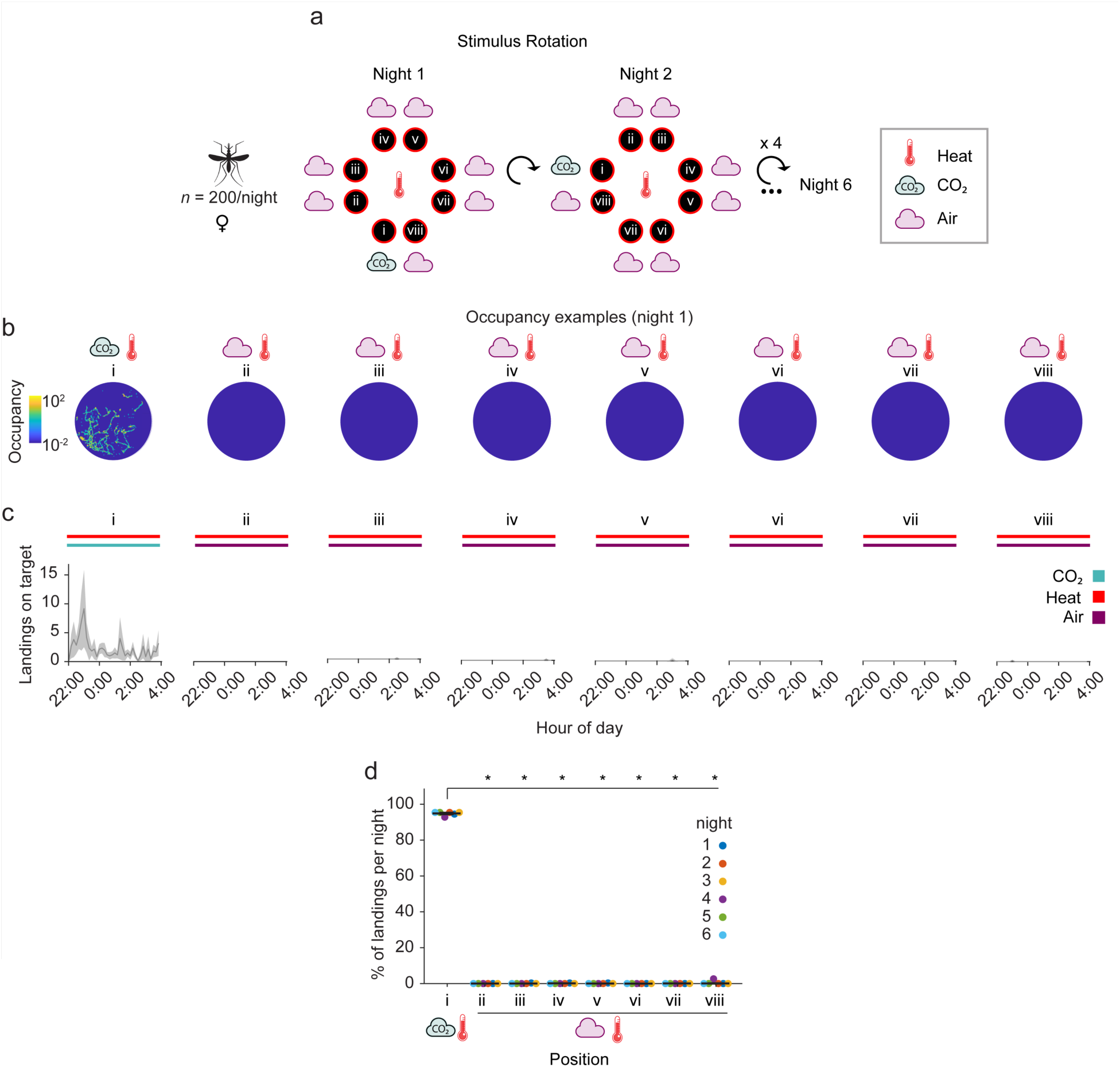
*Anopheles gambiae* prefers to land on heated targets baited with CO*_2_* over background air. (a) Stimulus position and rotation during 8-choice trials with these stimuli. The CO_2_ stimulus (position i) was rotated every night. The background air control positions (ii-viii) were labeled in reference to the CO_2_ stimulus position in a clockwise manner. All platforms were heated to 35°C throughout the experiment. (b) Heatmap of platform occupancy for an example night (night 1). (c) Number of mosquito landings on the OGTA platforms throughout the experiment. Mean ± SEM plotted. (d) Percentage of the total landings per platform per night. Mean ± SEM plotted. n = 6 nights. Significant differences from CO_2_: one-sided Wilcoxon signed rank test with Benjamini-Hochberg correction (* p < 0.05).

As the total number of landings varied every night (Fig. S2b), we subsequently calculated the relative attractiveness of each platform over replicate nightly trials (Fig. 3d). Overall, a strong and significant preference was observed for the heated platform baited with CO_2_, relative to the seven other heated platforms receiving only background air (Fig. 3d). Noticeably in this series of trials, wind appeared non-directional and wind speeds were very low (<0.2 m/s) (Fig. S2c), and we did not observe any cross contamination whereby CO_2_ appeared to obviously yield landings on adjacent platforms baited with background environmental air. We conclude that CO_2_ synergizes with heat to elicit *An. gambiae* landing on an artificial target mimicking human skin in a semi-field context, and heat alone is insufficient to evoke landings in this context.

### *Anopheles gambiae* prefers heated targets baited with human whole body odor relative to CO*_2_*

Volatile organic compounds (VOCs) present in human body odor have been found to synergize with CO_2_ to enhance attraction during mosquito host-seeking behavior ^23, 53, 57, 58^. Human whole body odor itself is a complex blend of hundreds of VOCs found in skin odor and breath, inclusive of ∼4% CO_2_ exhaled in human breath. To test olfactory preferences of female *An. gambiae* for human whole body odor relative to CO_2_ alone, we performed another eight-choice competition assay, whereby one heated OGTA platform was baited with human whole body odor from a sleeping human subject, and a CO_2_ stimulus that was 400ppm above background atmospheric concentrations was used to bait each of the seven other heated platforms (Fig. 4, Fig. S5 and Fig. S6).

**Figure 4.**
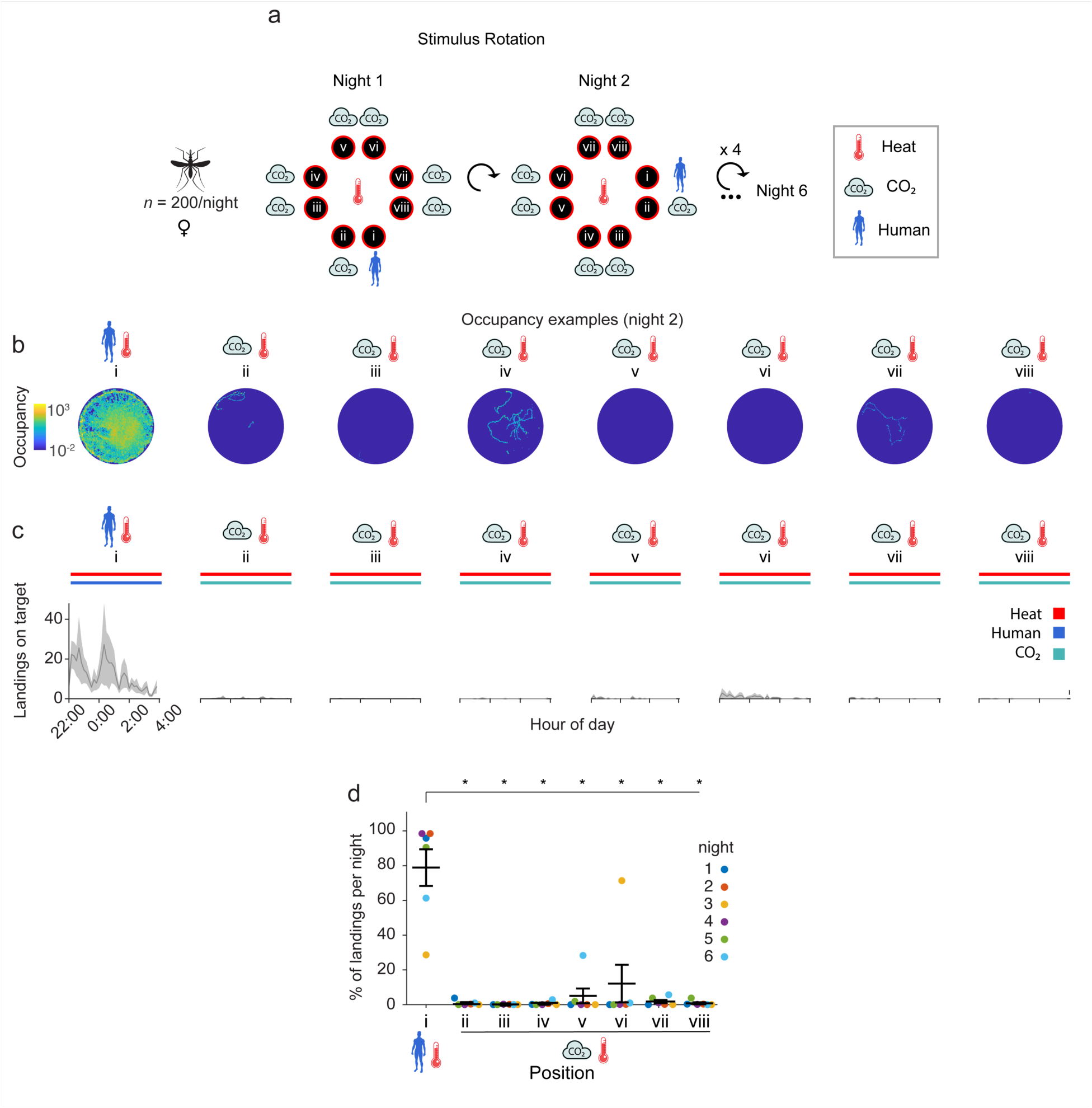
*Anopheles gambiae* prefers to land on heated targets baited with human whole body odor over CO*_2_*. (a) Stimulus position and rotation during 8-choice trials with these stimuli. The human stimulus (position i) was rotated every night. The remaining positions (ii-viii, CO_2_) were labeled in reference to the human position in a clockwise manner. All platforms were heated to 35°C throughout the experiment. (b) Heatmap of platform occupancy for an example night (night 2). (c) Number of mosquito landings on the OGTA platforms throughout the experiment. Mean ± SEM plotted. (d) Percentage of the total landings per platform per night. Mean ± SEM plotted. n = 6 nights. Significant differences from Human: one-sided Wilcoxon signed rank test with Benjamini-Hochberg correction (* p < 0.05).

In this multi-choice test, *An. gambiae* strongly preferred to land on the heated OGTA platform baited with human whole body odor relative to the other heated platforms that were baited with CO_2_ (Fig. 4b-c, Fig. S5b and Fig. S6). Across 6 replicate nights of these assays (Fig. 4a and Fig. S5a), the human whole body odor-baited heated platform was significantly preferred over the CO_2_-baited heated platforms (Fig. 4d and Fig. S5b). Interestingly, nights with lowest preference towards human odor were also nights with overall lower numbers of total landings (Fig. S5b), indicative of low mosquito activity. Similar to the previous trials, we determined that wind speed across replicate nights in the semi-field system during this series of trials was extremely low (Fig. S5c) and we found no clear indication of cross contamination, whereby any of the CO_2_ platforms immediately adjacent to the platform baited with human whole body odor had high levels of landing. These results demonstrate that *An. gambiae* strongly prefers 35°C targets baited with human whole body odor relative to CO_2_ alone.

### *Anopheles gambiae* prefers heated targets baited with the scent of one human over another

We next used the semi-field system in an eight-choice format to test whether female *An. gambiae* would exhibit olfactory preferences between two humans (Fig. 5, Fig. S7 and Fig. S8). In these assays performed over seven replicate nights, two heated OGTA platforms were baited with human whole body odor from two human subjects sleeping in separate tents, while the six other heated platforms were baited only with background air (Fig. 5a and Fig. S7a). During these assays, we observed that heated platforms that were only baited with background air received no landings (Fig. 5b-c and Fig. S7b). In contrast, high levels of landings were observed on both of the heated platforms baited with human whole body odor from either human.

**Figure 5.**
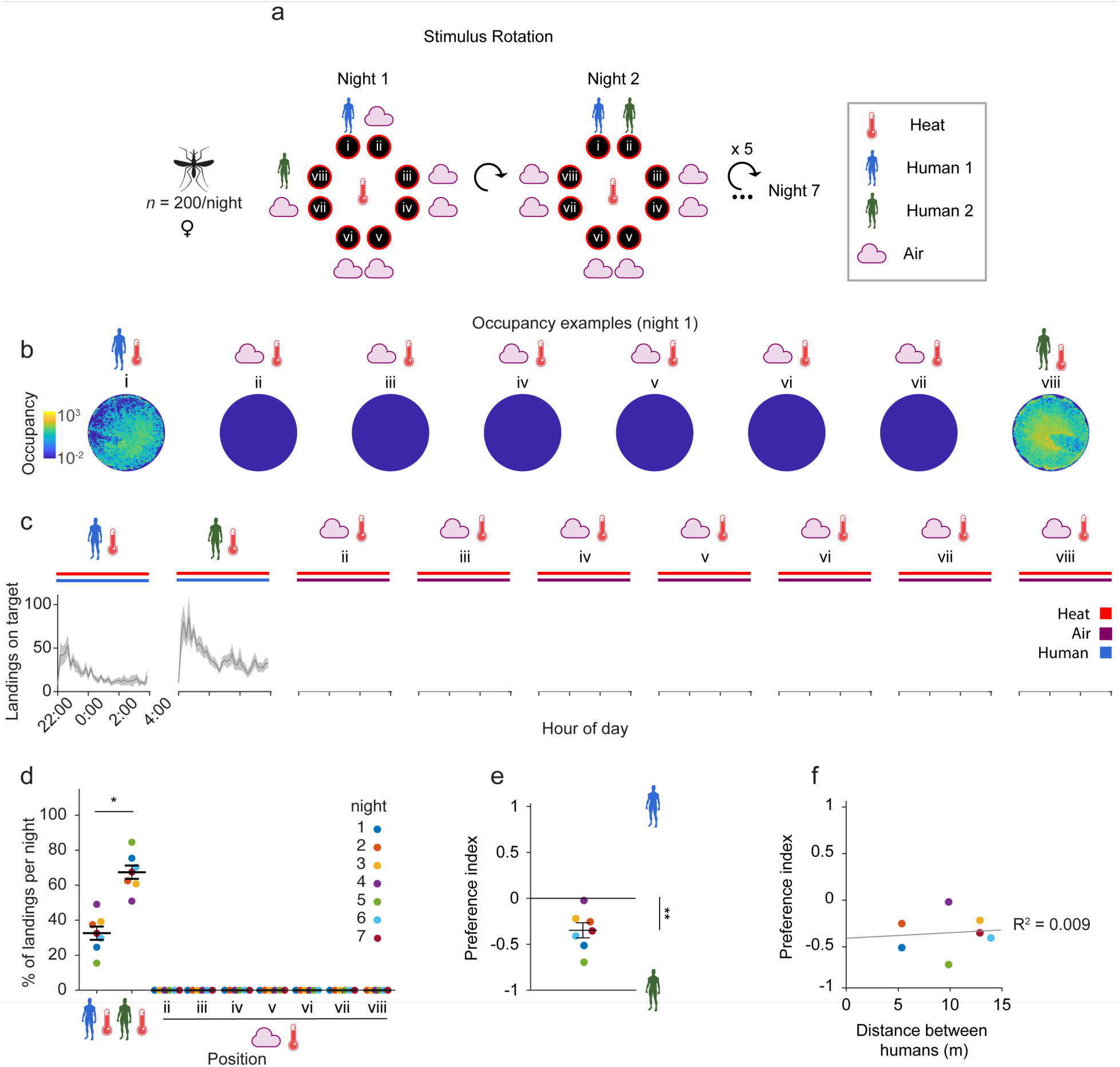
*Anopheles gambiae* prefers heated targets baited with the body odor of one human over another. (a) Stimulus position and rotation during 8-choice trials with whole body odor from two individual humans versus background air. The position of Human 1 (blue) and the position of Human 2 (green) relative to Human 1 were rotated every night. The control positions (air) and the position of Human 2 were labeled in reference to Human 1. Human 2 could take any of the positions ii-viii. (b) Heatmap of platform occupancy for an example night (night 1) where Human 2 took position viii. (c) Number of mosquito landings on the OGTA platforms. Mean ± SEM plotted. (d) Percentage of the total landings per platform per night. Mean ± SEM plotted. n = 7 nights (control positions have 6 data points since one of the positions was occupied by Human 2 every night). Significant differences between Human 1 and Human 2: two-sided Wilcoxon signed rank test (*p = 0.0156). Mean ± SEM plotted. (e) Preference index calculated from total landings on Human 1 and 2. One sample t-test for significant difference to 0: **p = 0.0054. (f) Effect of distance between Human 1 and 2 on the preference index. Coefficient of determination R^2^ = 0.009.

For both human subjects, we observed an initial peak in landing activity on the heated platforms within the first hour, followed by a sustained period of constant landing that was maintained throughout the assay period (Fig. 5c and Fig. S8). Noticeably, every single night of these trials, the heated platform that was baited with whole body odor from Human 2 received approximately double the number of landings relative to the heated platform baited with whole body odor from Human 1 (Fig. 5d and Fig. S7b). When considered in pairwise competition, a significant preference index for Human 2 over Human 1 was observed (Fig. 5e). Furthermore, this preference index was not influenced by the distance between the heated platforms baited with whole body odor from each of these human subjects (Fig. 5f); for instance, whether or not they were immediately adjacent to one another or farther away from each other.

In these trials, mean wind speed was low (Fig. S7c) and we saw no indication of cross contamination between platforms, whereby high levels of landings were recorded on heated platforms baited with background air which were immediately adjacent to a heated platform with human whole body odor. As rotating human positions in this eight-choice assay configuration every night had no effect on the tendency of *An. gambiae* to choose body odorants derived from Human 2 relative to those from Human 1, we conclude that this olfactory preference is likely due to differential attractiveness between whole body odor of these two humans, rather than position effects in the cage.

### *Anopheles gambiae* is differentially attracted to heated targets baited with the scent of certain individuals in a cohort of six humans

To extend use of this multichoice assay for *An. gambiae* olfactory preference to screen for differences in human attractiveness from larger groups of humans, we next tested a new cohort of six human subjects simultaneously in the semi-field system (Fig. 6, Fig. S9, Fig. S10 and Fig. S11). In this assay configuration over six replicate nights, six OGTAs were arrayed in a hexagonal configuration within the flight cage (Fig. 6a, Fig. S9a-b), yielding 15 pairwise comparisons between the individuals in this cohort per night. Relative to two-choice assay formats, use of this configuration therefore increases the throughput of screening to rank individuals by attractiveness by 15-fold.

**Figure 6.**
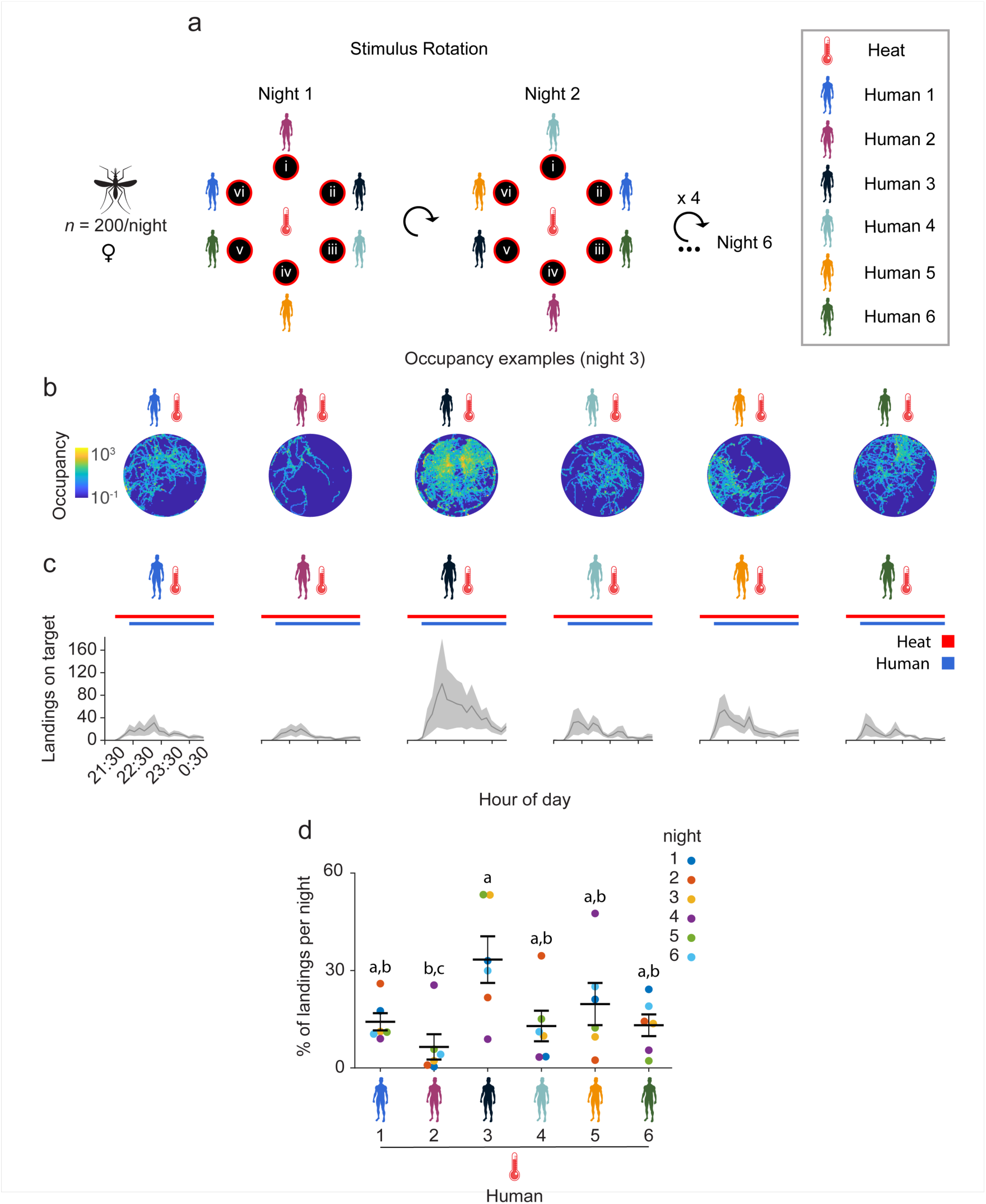
*Anopheles gambiae* differentially prefers body odor from certain individuals in a six-choice context. (a) Stimulus position and rotation during 6-choice trials with six humans. Humans were rotated randomly every night and could occupy any of the six positions (i-vi) numbered relative to their position in the cage. (b) Heatmaps of platform occupancy for an example night (night 3). (c) Number of mosquito landings on the platforms. Mean ± SEM plotted. (d) Percentage of the total landings per platform per night. Mean ± SEM plotted. n = 6 nights. Significant differences between humans: Fisher’s exact permutation tests with Benjamini-Hochberg correction (a-b: not significant, a-c: p = 0.033, b-c: not significant).

Since our previous experiments with two human subjects demonstrated that an olfactory preference is clearly observed during the first 3 hours of experimentation (Fig. 5c and Fig. S8), we recorded mosquito landing activity during both an acclimation period from 21:30 - 22:00 hours during which time heated platforms were only baited with background air from empty subject tents, and then from 22:00 -1:00 hours during which time the six humans occupied them. Consistent with the anthropophilic nature of *An. gambiae*, all six heated platforms baited with whole body odor from the human subjects yielded landings (Fig. 6b-c, Fig. S9c and Fig. S11). Importantly, no activity was observed during the initial 30-min acclimation period during which time the heated platforms were baited only with background air from the empty tents (Fig. 6c and Fig. S11), with landings starting as soon as whole body odor from each of the human was provided as an olfactory stimulus by humans entering the tents.

Across replicate nighttime trials, scent from Human 3 was observed to consistently be most attractive in this six-person cohort, on average having the highest number of landings and proportionally attracting the most mosquitoes on four out of the six nights tested (Fig. 6c-d, Fig. S9c and Fig. S11). In contrast, scent from Human 2 elicited very low landing activity and proportionally had the lowest attractiveness on four of the six nights, while the remaining subjects had similar intermediate levels of attraction (Fig. 6c-d, and Fig. S9c and Fig. S11). When ranked in terms of mean attractiveness, Human 3 was the most attractive, followed by Humans 5, 1, 6, 4, with Human 2 being the least attractive (Fig. 6d).

We quantified the percentage of landings per night per position and found that there was a slight, yet non-significant bias in the distribution of landings across different platforms within the flight cage arena (Fig. S10a), and consistent with previous competition trials, wind speeds were extremely low during these nighttime trials (Fig. S10b). To confirm that the differences in landings we observed were indeed due to differences in the attractiveness of whole body odor samples and not the position that they occupied relative to the cage, we simulated the likelihood of observing the same landing percentage differences between each pair of participants, under the null hypothesis that such biases arise only from position preferences (Fig. 6a, Fig. S9b and Fig. S10c). We estimated position preferences by averaging the nightly landing percentages at each location across experimental days. We then simulated 5,000 datasets by randomly reassigning landings to positions based on the average weighted position preferences and computed hypothetical landing percentage differences between individuals, averaged across experimental days. These simulations indicate that several pairs of humans have landing percentage differences that are highly unlikely to be observed solely due to a position bias (Fig. S10c), thus confirming that participant identity drives landing preferences. Further we discerned body weight across this six-person cohort did not correlate well with human attractiveness, suggesting that ratio-specific chemical constitution of human body odor, rather than overall body mass is the primary driver of *An. gambiae* olfactory preference for certain humans over others (Fig. S9d).

These data indicate that despite high levels of variability in the landing activity of *An. gambiae* (Fig. S9c and Fig. S11) and environmental conditions during nightly trials (Fig. S14 and Fig. S15), our multi-choice olfactory preference assay format is an efficient and powerful method to screen for humans at both ends of the attractiveness spectrum who are differentially attractive to *An. gambiae* relative to other individuals.

### Integrative use of whole body odor volatilomics identifies candidate volatile organic compounds modulating human attractiveness to *Anopheles gambiae*

To identify candidate chemical features of human scent modulating attractiveness to *An. gambiae*, we concurrently performed whole body volatilomics during the same six-human olfactory preference trials (Fig. 7, Fig. S12 and Fig. S13). To do this, we collected whole body odor from each human subject for chemical analysis by sampling air from tents onto thermal desorption tubes containing the chemical sorbent Tenax-TA. Air sampling from all subject tents occurred simultaneously during the first 45 minutes of human occupancy. Odor was collected onto the thermal desorption tubes by inserting them into a pump placed into the output ducting from each tent, after the ducting fan assembly (Fig. S9a). Samples were then analyzed by thermal desorption-gas chromatography/mass spectrometry (TD-GC/MS) to produce nightly chemical profiles of each human, resulting in a total of 36 human odor profiles (six individual human whole body odor signatures for six replicate nights) across the entire cohort throughout the study.

**Figure 7.**
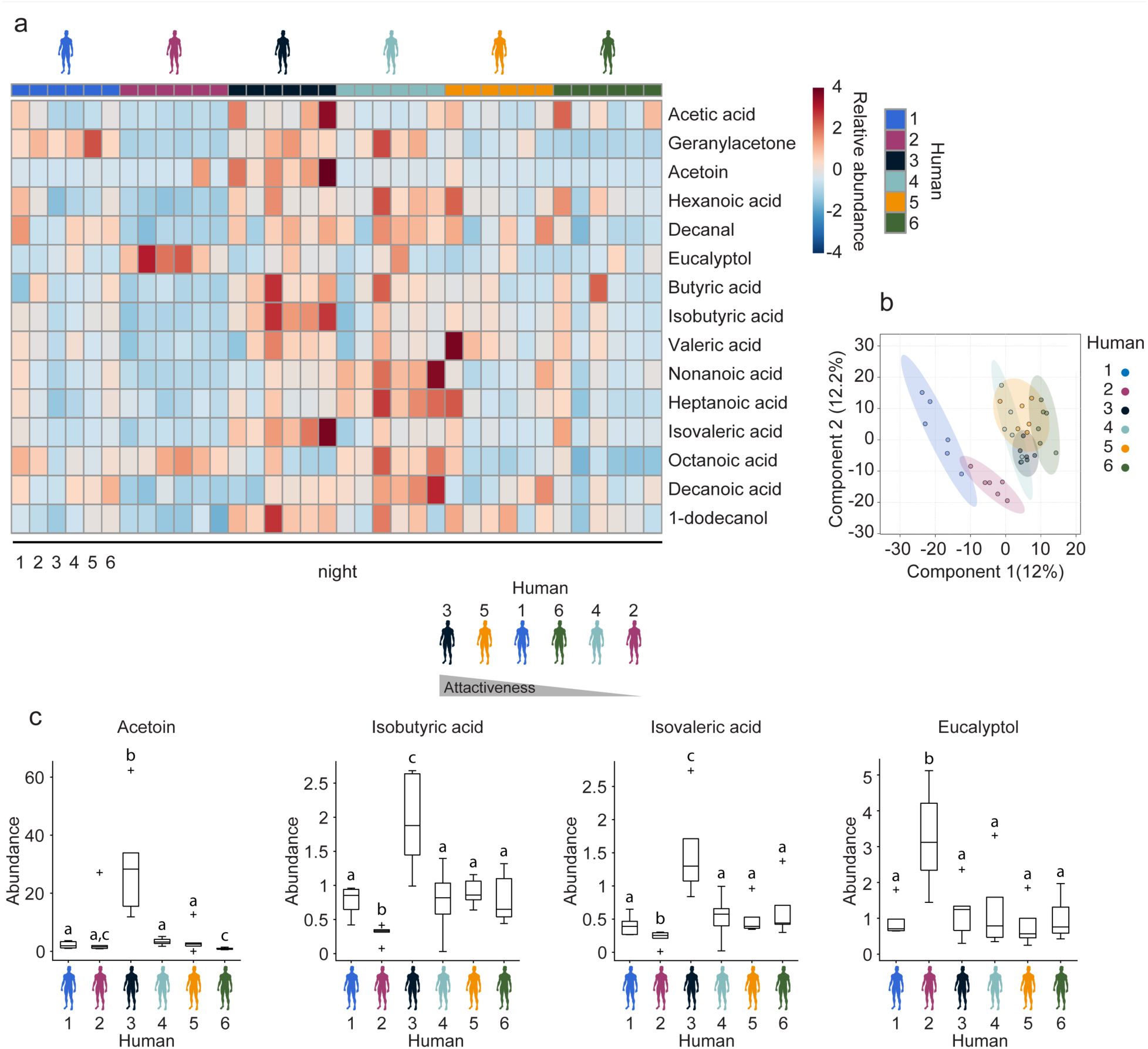
Whole body volatilomics reveals candidate compounds modulating *Anopheles gambiae* olfactory preferences for human scent. (a) Heatmap of 15 identified volatile organic compounds in human whole body odor demonstrating substantial variation between participants. Scale bar represents the amounts of analytes detected normalized to the internal standard, with red indicating a higher concentration and blue indicating lower concentration. Heatmap constructed using MetaboAnalyst 5.0. (b) PLS-DA score plot containing all chemical features detected. Ellipses indicate 95% confidence intervals. Plot constructed using MetaboAnalyst 5.0. (c) Variation in 4/15 compounds of interest: acetoin, isobutyric acid, isovaleric acid and eucalyptol detected in whole body odor from these humans. The line indicates the median, the box marks the lower and upper quartile, and the whiskers the 1.5 interquartile distance; outliers are indicated by black crosses. n = 6 nights. The letters indicate significant differences (p < 0.05) in compound abundance between humans: Fisher’s exact permutation tests with Benjamini-Hochberg correction.

In total, 1057 individual chemical features were detected across this dataset, predominantly consisting of human volatiles released via the skin and breath of participants, exogenous contaminants from the tents and ducting, and volatiles introduced into the tents from the external environment, as well as features of unknown chemical identity due to limitations in current mass spectral libraries and standards for identification. All features in this dataset were initially used to generate a partial least squares discriminant analysis (PLS-DA) model (Fig. 7b) to visualize variation between samples from each human subject and aid in the identification of features demonstrating significant variation.

We next leveraged our previous analyses of human whole body headspace ^48^ to target the primary chemical constituents of human whole body odor, confirming the identity of 40 known VOCs detected in whole body odor in this semi-field dataset (Fig. S12). Air-borne human odorants from numerous chemical classes were detected across subjects, including aldehydes, ketones, alcohols, carboxylic acids, and hydrocarbons (Fig. 7a and Fig. S12). Of these identified compounds, 15 were found to exhibit substantial variation in the whole body emissions across the human cohort (Fig. 7c and Fig. S13) including acetic acid, butyric acid, isobutyric acid, valeric acid, isovaleric acid, hexanoic acid, heptanoic acid, octanoic acid, nonanoic acid, decanoic acid, acetoin, geranylacetone, decanal, 1-dodecanol, and eucalyptol. These compounds were conserved in the skin and breath emissions of all participants (Fig. 7a and Fig. S12) but were released at different rates.

The majority of identified compounds differing between participants were carboxylic acids, which were generally more abundant as a class of molecules in the scent of Human 3 (the most attractive human subject) and least abundant in the scent of Human 2 (the least attractive human subject) (Fig. 7c and Fig. S13). Specifically, Human 3 exhibited significantly higher abundances of the carboxylic acids: butyric acid, isobutyric acid, isovaleric acid, as well as the skin microbe-generated volatile acetoin (3-hydroxy-2-butanone) relative to all other human subjects (Fig. 7c). Conversely, Human 2 exhibited lower abundances of the vast majority of these 15 VOCs (Fig. 7c and Fig. S13), including significantly lower abundances of isobutyric acid and isovaleric acid (Fig. S13). A notable exception was the monoterpenoid eucalyptol which was significantly elevated by 2-fold in abundance in this individual relative to all others (Fig. 7c).

These data indicate that despite nightly variation in chemical signatures of whole body odor between human subjects, a number of volatile organic compounds were consistently released at greater or lower concentrations by individuals demonstrated to be at either end of the attractiveness spectrum in this targeted cohort of six humans. Integrative use of this multi-choice olfactory preference assay with whole body volatilomics therefore has the potential to yield candidate chemical modulators of human attractiveness to *An. gambiae*.

## Discussion

The African malaria mosquito is a highly anthropophilic mosquito species that blood feeds preferentially on humans ^59–62^, exhibiting an innate olfactory preference to seek out human body odor. In addition to displaying strong inter-specific host preference for the scent of humans over other vertebrate host animals ^63^, *An. gambiae* has also been observed to consistently exhibit intra-specific host preferences to seek out and blood feed on certain humans over other individuals ^3–8^. The chemosensory and other biological factors driving anthropophilic host preferences and inter-individual differences in human attractiveness to *An. gambiae* and other malaria vectors remain poorly understood. New mosquito behavioral paradigms are therefore needed to address these important questions of both fundamental and applied significance to combat malaria and other mosquito-borne diseases globally.

The large-scale, semi-field system we have developed at Macha, Zambia described here within provides a contained, yet naturalistic setting to perform mosquito multi-choice olfactory preference assays with optimized machine vision analyses. Relative to complementary methods for quantifying mosquito host preference ^5, 13, 18, 20, 37, 42–44^, our semi-field system provides an exposure-free method to quantify mosquito host preference from multiple individuals at a time during peak periods of *An. gambiae* host-seeking activity ^47^ in the hours flanking midnight. Using multiple Odor-Guided Thermotaxis Assays (OGTAs) arrayed in competition, we observed that female *An. gambiae* display a hierarchical olfactory preference to land on heated OGTA platforms mimicking human skin temperature (35°C), when they are baited with carbon dioxide (CO_2_) over background atmospheric air, human body odor over CO_2_, and the scent of one human over another.

Interestingly, we observed that when heated OGTA platforms were only baited with background air, the visual and physical stimuli from these basal assay components were insufficient to evoke landings by *An. gambiae* in this large flight cage arena. These results coupled with the hierarchical nature of behavioral preferences shown by *An. gambiae* to olfactory stimuli of increasing chemical complexity, suggest that at enhanced spatial scales, multi-sensory integration between visual cues, heat, CO_2_ and other human body odorants ^23, 46, 53–55^ act critically to bolster the fidelity of host localization by this disease vector.

The human body emits hundreds of volatile organic compounds (VOCs) in exhaled breath and skin emissions ^64^, and our prior research has demonstrated considerable heterogeneity in the chemical complexity of whole body emissions between humans ^48^. This natural variation in the types and abundances of VOCs emitted by the human body has long been hypothesized as a key factor driving mosquito host preference ^65^. In this study, we integrated whole body volatilomics with our multi-choice olfactory preference assay to provide insights into candidate air-borne VOCs modulating human attractiveness to *An. gambiae*.

Our chemical analysis of whole body headspace in a target cohort of six humans exploited thermal desorption-GC/MS (TD-GC/MS) to specifically target compounds solely present in human headspace, thus only evaluating components sufficiently volatile to travel from the tent housing each human through the attached ducting path towards the flight cage arena. We chose this specific analytical method for initial application with whole body volatilomics, in an attempt to identify human-derived VOCs that act distally to attract *An. gambiae* towards humans. A large number of previous studies profiling human-derived volatiles as candidate mosquito attractants or repellents commonly use contact-based sampling methods biased towards less volatile compounds, such as glass beads rubbed across the hands or feet ^22, 66, 67^ or absorbent materials pressed against the skin ^37, 68^ that are subsequently extracted for analysis; in contrast likely representing important classes of molecules for close-range olfactory and gustatory attraction.

Strikingly, we discerned that the majority of human body odorants found to significantly differ in abundance from the scent of humans at both end of the attractiveness spectrum using whole body volatilomics were carboxylic acids. Specifically, we discerned that butyric, isobutryric and isovaleric acids were significantly enriched in the most attractive subject, while these latter two carboxylic acids were significantly diminished alongside several other carboxylic acids in the least attractive of humans from this six-person cohort. Introduced into human odor both via the sebaceous glands on the skin and by skin microbiota ^69^, carboxylic acids have been a compound class of great interest in the study of the chemical basis of mosquito host-seeking for decades ^33^. Numerous studies have demonstrated the physiological and behavioral activity of carboxylic acids in both *An. gambiae* and *Ae. aegypti*. Short- and medium-chain carboxylic acids have been found to elicit physiological responses in both electroantennogram (EAG) and single sensillum recordings (SSR) in *An. gambiae* mosquitoes ^70, 71^. Smallegange *et al.* found that a mixture of 12 carboxylic acids synergized with ammonia and lactic acid to elicit *An. gambiae* attraction in a dual-port olfactometer assay ^72^. Furthermore, a dual-port oflactometer-based screen ranking the attractiveness of foot odorants captured onto glass beads from 48-humans revealed that lactic acid, 2-methylbutanoic acid and tetradecanoic acid were highly enriched in abundance amongst individuals classified as being highly attractive to *An. gambiae* ^22^.

More recently in a study examining the attractiveness of arm odor from different humans to *Ae. aegypti*, De Obaldia *et al.* found using contact-based sampling that highly attractive individuals had higher abundances of a number of medium and long-chain carboxylic acids (C_10_-C_20_) ^37^. We observed similar trends in our study using air sampling, with the exception that we detected more highly volatile carboxylic acids (C_2_-C_10_), with these being more or less abundant in whole body odor of human subjects at both ends of the attractiveness spectrum. It is therefore tempting to speculate that abundance of certain carboxylic acids may be both important close-range and distal olfactory cues that potently modulate human attractiveness across these two anthropophilic mosquito species.

We also observed that a skin-microbe generated compound acetoin (3-hydroxy-2-butanone) was highly enriched in the most attractive human subject in the six-choice trial. Acetoin is amongst the most abundant of VOCs detected in the human whole body volatilome ^48^. The bacteria *Staphylococcus epidermidis* and *Staphylococcus aureus* have been demonstrated to produce acetoin, both of which are abundantly present on human skin ^14, 73^. The role of acetoin in mosquito host-seeking has not been well studied, although some evidence does suggest this compound may be implicated in *An. gambiae* attraction to humans. For instance, in a screen of microbial volatiles as mosquito attractants, Verhulst *et al* found acetoin increased the efficiency of mosquito traps when tested with *An. gambiae*, although this was highly dependent on acetoin concentration ^74^. An important area for future research will thus be to leverage our semi-field system with expanded cohorts of humans to understand the influence of the human skin microbiome ^75, 76^ on inter-individual variability in the production of acetoin and the other carboxylic acids in modulating mosquito behavior. In the future, reconstitution of synthetic blends containing these components and other conserved human odorants, may be further used to test their sufficiency to evoke mosquito olfactory attraction in combination with the OGTA assay. Such studies may have the potential to yield blends that are super attractive to *Anopheles gambiae* and other anthropophilic disease vectors for enhanced vector surveillance and control, by mimicking the chemistry of scent signatures from highly attractive humans.

Conversely, just as the presence of specific attractant cues in human odor may lead to an increase in human attractiveness to mosquitoes, the presence of repellent volatile compounds may have the opposite effect. Interestingly, the monoterpernoid eucalyptol (1,8-Cineole) was highly abundant in the body odor of the least preferred human subject in the six-person cohort. Eucalyptol is a molecule with known mosquito repellency ^77–79^ and deodorizing action ^80^ that has been frequently detected in the skin and breath emissions of humans ^64^. This compound is likely derived from plant-based foods in diets or exogenous products (European Comission, 2002). Expansion of the behavioral screens described here to identify individuals whose scent is consistently not attractive to mosquitoes when placed in competition with others, may therefore have the potential to reveal both established and novel biomarkers of human repellency to mosquitoes.

Application of air- and contact-based sampling methods in combination may also yield high-content information about features of the human scent signature that modulate mosquito attractiveness to humans over a range of spatial scales, and further assist in the discovery of novel mosquito attractants and repellents. Future studies may also benefit from using additional TD-GC/MS sorbent materials to those described here to expand the types of human body emissions analyzed, expanding discovery of airborne molecules modulating mosquito behavior into different parts of chemical space. Similarly, the use of alternative analytical techniques could further increase the types of compounds detected. Direct MS techniques, such as proton-transfer reaction MS (PTR-MS) and gas-specific monitors ^82^ could be used for the real-time analysis of compounds that are not readily captured by sorbent materials including compounds such as ammonia and CO_2_.

In summary, we have developed and validated a large-scale, semi-field system in Zambia to quantify mosquito olfactory preference for whole body odor sourced from different humans, and identify candidate chemical features modulating human attractiveness to *Anopheles gambiae*. Given the enhanced throughput and comparative power of our multi-choice assay, with potential to compare the attractiveness of up to eight humans at a time, we propose this system will be a highly useful resource to rapidly screen large cohorts of humans to identify chemical correlates of mosquito attractiveness to humans in *Anopheles gambiae* and related malaria vectors. From an applied perspective, the OGTA we describe may additionally be used as an exposure-free method to bench mark the comparative behavioral efficacy of next-generation mosquito repellents and synthetic blends that aim to faithfully mimic human scent. By extension, we propose this semi-field system could readily be used to study how factors such as diet, pregnancy, the human microbiome and infectious state drive preferential attraction of mosquitoes to humans in a standardized fashion under naturalistic conditions.

## Materials and Methods

### Mosquito Stock Maintenance

Laboratory experiments at Johns Hopkins Malaria Research Institute (JHMRI) were performed with the *Anopheles gambiae Keele* strain. Mosquitoes were maintained with a 12hr light:dark photoperiod at 27°C and 80% relative humidity using a standardized rearing protocol. Adults were provided with a 10% sucrose solution for colony maintenance.

Semi-field system experiments at Macha Research Trust, Zambia (MRT) were performed with the *Anopheles gambiae Kisumu* strain. Mosquitoes were laboratory reared in an insectary facility adjacent to the flight cage. Mosquitoes were maintained with a natural light:dark cycle at 26-30°C and 70-80% relative humidity using a standardized rearing protocol. Adults were provided with a 6% glucose solution for colony maintenance.

### Odor guided thermotaxis assay

An odor guided thermotaxis assay (OGTA) was engineered in a 30cm x 30cm x 30cm cage (Bugdorm-1 Insect Rearing Cage, MegaView Science Co.). The assay consisted of a heated platform placed on the floor of the cage, made from a black aluminum disk of 10cm in diameter with a Peltier element (Aideepen TEC1-12706) glued at the bottom. The Peltier element was controlled by an Arduino UNO microcontroller and a thermistor (WM222C, Sensor Scientific, Inc.) to keep the temperature at 35°C, mimicking human skin temperature. Surrounding the platform was a 3D printed ring with 12 infrared LEDs (840nm) powered by the Arduino to illuminate mosquitoes landing on the platform. A Raspberry Pi camera module (Camera Module 2 Pi NoIR) was placed on top of the cage to record mosquito activity and all recordings were carried out at 10fps. Five percent CO_2_ or clean synthetic air (Airgas) were brought into the cage with 1/4” vinyl tubing (Cole Parmer 06405-02), that was placed immediately adjacent to the platform in a set position in the 3D printed ring, at a flow rate of 2mL/s. The end of the tubing was capped with mesh to prevent mosquitoes from flying into it.

### Odor guided thermotaxis assay under laboratory conditions

Twenty-five mated and nulliparous *Anopheles gambiae* Keele strain females (4-7 day old) were used for each replicate experiment at JHMRI to initially pilot the OGTA configuration prior to its adaptation for semi-field use. Approximately 30 hours prior to experimentation, mosquitoes were cold anesthetized at 4°C, and placed in a 3D-printed release trap, and provided with a 10% sucrose solution on a saturated wet cotton ball overnight. The sucrose solution was removed and replaced by a cotton ball soaked in dH_2_O the next day, 12 hours prior to start of each experiment, to encourage host-seeking behavior.

Three or four hours before the experiment, the release trap was placed in a fitted receptacle on the front of OGTA cage and the assay placed in an isolated room maintained at 27°C and 60% humidity. In experiments using heat, the landing platform was warmed to 35°C to simulate human skin temperature. The platform was left unheated in no-heat experiments, but the temperature was monitored over the duration of the assay using the thermistor. For all experimental replicates, a paper towel wick was inserted into a round glass bottle (Fisherbrand FB02911944) containing dH_2_O that was placed in each assay to prevent mosquito desiccation throughout the night.

To initiate each experiment, the release trap was opened, and mosquitoes were allowed to freely move into the enclosure. All lights in the room were turned off except for a night light (MAZ-TEK YOOUS, 25 lumens) placed on the floor in front of the OGTA to simulate nighttime conditions. Mosquitoes were allowed to acclimate for 3-4 hours before the start of the experiment to ensure that enough time passed since an experimenter was in the room, minimizing the possibility of human odors influencing mosquito behavior. At 20:00 hours mosquito landings on the platform were recorded for 30 minutes at 10fps. Five percent CO_2_ or clean synthetic air was pulsed using a solenoid valve (Clippard ETO-3-12) and valve controller (Automate Scientific ValveLink 8.2) into the assay at the set position immediately adjacent to the platform at 20:00 hours and 20:15 hours for 1 minute each at a flow rate of 2 mL/s. Mosquito landings were tracked using ivTrace (Jens P. Lindemann, Bielefeld University). All the trajectories were manually corrected and then analyzed with MATLAB (The Mathworks Inc., R2018b).

### Semi-field system

Semi-field experiments were performed in a custom flight cage facility (20m L x 20m W x 2.5m H) constructed at Macha Research Trust, Chroma District, Southern Province, Zambia. Eight one-person tents (one-person Swag tent, #8101, Kodiak Canvas) were distributed evenly around exterior walls of the screened flight cage (two tents per side) to accommodate sleeping humans and other olfactory stimuli such as CO_2_ that were introduced into them. For participant comfort and protection from inclement weather, tents were erected on stretchers (Outfitter XXL Camp Cot, #120A, Teton Sports) that were placed on dedicated concrete slabs, each 15m away from the perimeter of the central flight cage. The flight cage was positioned in a large open field on a gentle slope flanked at its margins by native foliage and grasses. A corrugated metal roof was installed on this facility to protect the central screened flight cage arena and enclosed assay components from weather-related damage due to the heavy seasonal rainfalls encountered in the Macha region during the wet season, which typically spans from November-April in Southern Zambia. An anteroom to facilitate secure entry and exit from the central flight cage arena was positioned on the northeastern side of the cage. To ensure that aluminum ducting piping olfactory stimuli from each tent into the flight cage were of uniform length and spatial position, each concrete tent slab was made to be level with the slab of the central flight cage arena. In a clockwise manner from the northeastern side of the cage, the tops of each pair of side tent slabs were 27cm and 23cm; 74cm and 95cm; 125cm and 120cm; and 30 cm and 8cm from the ground, respectively.

Each tent was modified using a ducting flange so that 4” air conditioning aluminum ducting could be connected to the front canvas surface of the foot of the tent. The tent was then connected to an inline fan (Inline Blower, Rule) powered by a 12V battery (12AH). Eight window access ports on the sides of the flight cage arena (two per side) allowed the 4” aluminum ducting from the tent fans to be connected to the cage. Inside the cage, each window access port was connected via ducting connectors to a modified version of the OGTA described above. The opening of the 4” ducting was secured in place in front of the platform using a U-bracket, and it was covered with aluminum screen to prevent mosquitoes from flying into it. The fan was used to duct air from the tents onto the OGTAs at a speed of 0.8 m/s. Airflow at the output of the ducting on the OGTA was calibrated using a handheld anemometer (Testo 405i) and airflow adjusted by modifying fan speed using a pulse width modulator (RioRand RR-6-90V-15A-PSC). The mosquito landing platform was warmed to 35°C and illuminated with 12 infrared LEDs controlled by an Arduino, and mosquito landings were recorded with the video camera placed directly over the platform. The Raspberry Pi and Peltier element were powered by two 12V batteries, and all electronic components were placed in a black container that had the landing platform and infrared sensitive camera (Raspberry Pi Camera Module 2 NoIR) held by a camera stand (Myrin Korea B00AUJHKC2) on top of the container lid. Four water containers were placed in an evenly spaced configuration in the corners and two wet towels were placed in the center of the flight cage to increase local humidity around the OGTAs during assays.

### Mosquito preparation for semi-field behavioral experiments

To prepare for each evening of assays in the semi-field system, 200 mated and nulliparous *Anopheles gambiae* Kisumu strain females (laboratory reared and 3-5 day old) were aspirated into a 30 x 30 x 30 cm release cage (Bugdorm-1 Insect Rearing Cage, MegaView Science Co.) at 8:00 hours and glucose starved for 12 hours prior the start of experimentation. During glucose deprivation, mosquitoes were provided with wet cotton balls soaked with dH_2_O as a water source and maintained under insectary conditions with natural light conditions until the evening. Mosquitoes were released into the center of the flight cage at 20:00 hours and allowed to acclimate to semi-field conditions for 2 hours. Landing platforms were heated to 35°C and infrared LEDs on all platforms were switched on immediately before mosquito release at 20:00 hours and remained on for the entirety of the experiment. Mosquito activity on each platform was recorded at 10fps for the duration of the experiments, which occurred with varying lengths of duration depending on the experimental configuration between 22:00 hours and 4:00 hours (see below). Mosquitoes were collected and totally cleared from the cage every morning using mechanical aspiration (Prokopack 1419, John W. Hock Company).

### Preference for CO*_2_* versus environmental air

To quantify *An. gambiae* olfactory preference for warmed targets baited with CO_2_ relative to environmental air, eight OGTAs were arrayed in an octagonal pattern in the center of the flight cage and heated to 35°C. A 100% CO_2_ source was then introduced into a single tent using 1/4” tubing at a flowrate of 300mL/min from a compressed tank using a tank regulator and flow meter for the entire duration of the experiment, yielding an ∼ 0.08% CO_2_ stimulus at the output of the ducting onto the OGTA after mixing with environmental air being drawn through the tent by the ducting fan assembly at rate of 0.8 m/s (i.e. ∼ 400 ppm above background atmospheric CO_2_ concentrations). CO_2_ concentration at the output of OGTA ducting using these flow parameters was determined under simulated assay conditions in the laboratory using an identical tent configuration and ducting flow rate using a CO_2_ meter (IAQ Mini, CO_2_Meter, USA). No stimulus was added to the seven remaining tents, and air from all tents was brought into the cage and onto the platforms at a speed of 0.8 m/s as described above. The CO_2_ stimulus was inserted into a different tent every night to control for possible position effects and experiments were carried out for 6 consecutive nights. Recordings were carried out for 6 hours from stimulus onset at 22:00 hours to 4:00 hours. Mosquito landings were tracked using ivTrace (Jens P. Lindemann, Bielefeld University). All the trajectories were manually corrected and then analyzed with MATLAB (The Mathworks Inc., R2018b).

### Study Participants

For one-person experiments, one healthy adult male was used, while for two-person experiments, an additional healthy adult male was employed. For six-person experiments, a new cohort of six healthy adults was recruited consisting of two females and four males. Study participants in the two-person and six-person experiments were assigned random numerical identifiers. Participants in the six-person cohort were provided with odor-free shampoo and body wash (Vanicream, USA) to wash with each day of the study prior to sampling. After washing, participants were requested not to use any other cleaning products, deodorants, fragrances or cosmetics. For 12 hours prior to each experiment, participants were also asked to refrain from the consumption of alcohol and odorous foods such as onions and garlic. Prior to entering the tents, participants changed into a set of scrubs (65/35 polyester/cotton, SmartScrubs, Phoenix, USA) washed only in water to reduce the introduction of exogenous VOCs from non-standardized clothing. Clothing was not standardized during the one-person and two-person experiments as these occurred during an early pilot phase of assay development, prior to later recruitment of the six-person cohort after standardization of assay conditions. This study was approved by the Johns Hopkins Bloomberg School of Public Health (JHSPH) Institutional Review Board (IRB no. 00018691), Macha Research Trust (MRT) Institutional Review Board (IRB no. E.2021.06) and the National Health Research Authority of Zambia (NHRA0000004/08/04/2022).

### Preference for human whole body odor versus CO*_2_*

To quantify *An. gambiae* olfactory preference for warmed targets baited with human whole body odor relative to CO_2_, a total of eight OGTAs heated to 35°C and associated tents were used. One male human subject slept in one tent over consecutive nights to provide a replicate human whole body odor stimulus. A 100% CO_2_ source from a compressed tank was split equally to achieve a flow rate of 300mL/min into each of the seven other tents, yielding an ∼0.08% CO_2_ stimulus at the output of the ducting of each concordant OGTA (i.e. ∼ 400 ppm above background atmospheric CO_2_ concentrations) after mixing with environmental air being drawn through each tent by the ducting fan assemblies. Air from the tents was brought into the cage and onto the platforms at a speed of 0.8 m/s as described above. To avoid position effects and prevent possible bias due to contamination of the tent and landing platform by human scent, the human subject together with the tent, ducting, and platform were rotated every night. This was to ensure the human would sleep in the same tent, but at different positions relative to the cage. The same person was tested over 6 six consecutive nights. Recordings were carried out from stimulus onset at 22:00 hours to 4:00 hours. Mosquito landings were tracked using ivTrace (Jens P. Lindemann, Bielefeld University). All the trajectories were manually corrected and then analyzed with MATLAB (The Mathworks Inc., R2018b).

### Preference between two humans

To quantify *An. gambiae* olfactory preference for human whole body odor from two different humans, an assay configuration with 8 OGTAs and 8 tents were used. Two male human subjects were placed in individual tents and no stimulus was added to the remaining tents. All eight platforms were heated to 35°C. Air from the tents was brought into the cage and onto all eight platforms at a speed of 0.8 m/s as described above. To avoid position bias, the position of the subjects was rotated relative to the cage as described for the human versus CO_2_ experiments. Additionally, the position of human 2 relative to human 1 was changed every night until all possible combinations were tested over 7 consecutive nights. The same two persons were tested every night. Recordings were carried out for 6 hours from the moment the subjects entered the tent at 22:00 hours to 4:00 hours. Mosquito landings were tracked using ivTrace (Jens P. Lindemann, Bielefeld University). All the trajectories were manually corrected and analyzed with MATLAB (The Mathworks Inc., R2018b).

### Preference between six humans

To quantify *An. gambiae* olfactory preference for human whole body odor sourced from 6 different humans, an assay configuration with six OGTAs and six tents was used. In these tests, OGTAs were arranged in a hexagonal configuration with each unit spaced equidistantly 5.3 m from the two neighboring units. All OGTAs were heated to 35°C and air from the tents was brought into the cage and onto the platforms at a speed of 0.8 m/s as described above. To avoid position bias, the position of the subjects was randomly rotated every night relative to the cage. Each subject slept in the same tent every night, but unlike previous experiments the platforms and ducting were maintained in the same position. Recordings started 21:30 hours to record landings in absence of a human stimulus. Humans were introduced at 22:00 hours and recordings carried on until 1:00 hours for a total of 3.5 h. The same cohort of six subjects was tested over 6 replicate nights.

Given the high level of overall mosquito activity observed in six-human choice experiments, trajectories were calculated using a custom-made program on Python (Python 3.8.13). Mosquito landing trajectories were reconstructed using a background-subtraction approach. Recording backgrounds were determined based on rolling windows of 5-minute (3000 frames) duration, with background intensities defined as the 15th percentile with respect to the time axis of each window. A background-subtracted pixel value (p_i,j <in> [0, 255]) was considered a region of interest relative to its corresponding background intensity (b_i,j) if p_i,j >= 1.25 * b_i,j and p_i,j - b_i,j >= 25. Following erosion (2 iterations) and dilation (24 iterations), these regions of interest were stored in an intermediate MicroFlyMovieFormat representation to facilitate trajectory reconstruction and manual verification.

Mosquito landing trajectories were determined by linking previously identified bright pixel clusters across frames based on the euclidean distance between their centroid locations, with closer matches taking precedence. A maximum linking distance of 50 pixels was found to robustly identify trajectories. Trajectory frame ranges were considered to represent a landing if their velocity fell below 20 pixels/frame (“low velocity”), or if a timepoint occurred within a <=5-second window between two low-velocity periods. To ensure no false positive trajectory detections occur due to background artifacts, trajectories with no significant displacement (<=20 pixels span) were rejected. Furthermore, due to the occurrence of flickering of the infrared illumination on some recording days, we excluded putative trajectories that remained in a 20-pixel vicinity of bright background artifacts corresponding to areas of high illumination surrounding the LEDs. Maps of such light artifacts were defined as pixel locations whose 95th percentile (taken across frames) exceeded the overall 90th percentile of pixel intensities in all frames and positions. All the trajectories were processed and analyzed with MATLAB (The Mathworks Inc., R2018b).

### Human whole body volatilomics

Whole body VOC samples were collected using low-power pumps (Pocket Pump, SKC Inc., USA) with attached sorbent tubes that were positioned inside the 4” ducting transferring human odor from the tents. To facilitate sampling of air from the ducting in a closed fashion, pumps with attached sorbent tubes were placed within the sealed arm of a 4” three-way ducting splitter that was positioned 1m from the base of the tent downstream of the fan assembly. Whole body odor samples were collected directly onto Tenax-TA thermal desorption (TD) tubes (#020810-005-00, 6 x 60 mm, Gerstel, USA) at 250 mL/min^-1^ for 45 minutes. Prior to sampling, TD tubes were conditioned at 300°C for 60 min in a stream of nitrogen at 50 mL/min^-1^ using the Gerstel tube conditioner (TC2, Gerstel, USA) and then spiked with 40 ng of 4-methylphenol (99+%, TCI America, USA) as an internal standard. Whole body VOC sampling commenced at 22:00 hours each night upon entry of participants into the tents.

Samples were analyzed using thermal desorption gas chromatography/mass spectrometry (7890B GC, 5977N MSD, Agilent, USA). Tenax-TA tubes were placed in a Gerstel Thermal Desorption Unit mounted onto a Gerstel Cooled Injector System (CIS4) PTV inlet (Gerstel, USA). Analytes were desorbed in splitless mode starting at 30°C followed by an increase of 720°C /min to 280°C with a 3 min hold. Analytes were then transferred into the inlet which was held at -70°C and then heated at 720°C/min to desorb analytes onto a HP-INNOWAX capillary column (30 m length x 0.25 mm diameter x 0.25 µm film thickness). The GC oven was programmed with an initial temperature of 40°C with a 2 min hold followed by an increase of 6°C /min to 250°C and held for 5 mins. A helium carrier gas with a flow rate of 1.2 mL/min^-1^ was used. The MS analyzer acquired over a range of m/z 30-300 and was operated in EI mode. The transfer line and ion source were set to 250°C and 230°C respectively.

Raw data were converted to mzML formats using MSConvert version 3.0 and processed in MSDIAL version 4.90 for peak alignment, integration and deconvolution. MSDIAL parameters were as follows: average peak width 20 scans, smoothing level 3 scans, minimum peak height 1000, deconvolution sigma window 0.5, and EI similarity cutoff 70%. Chromatographic peaks occurring above the limit of detection were normalized to the internal standard. Identification of compounds was achieved by comparison of mass spectra with the NIST Mass Spectral Library version 2.2 and the MassBank of North America spectral libraries and retention time matching with analytical reference standards. MetaboAnalyst 5.0 was used for the production of heat maps and chemometric analysis.

### Statistical Analyses

Wilcoxon signed-rank and Fisher’s exact permutation tests were used to quantify statistical differences in landing percentages and total landings during single and multiple comparisons, respectively. A one-sample Kolmogorov-Smirnov test was used to test for data normality. Statistical differences in compound abundance were calculated using Fisher’s exact permutation tests. The Benjamini-Hochberg procedure was used to correct for multiple comparisons. All statistical analyses were performed with Matlab (The Mathworks Inc., R2018b). The p-values for every comparison in the study can be found on Data Sheet S1.

For six-person experiments, to evaluate the statistical significance of individual landing preferences considering the potential for overall preference bias by physical position in the flight cage arena, we used the following randomization test approach. For each position, a landing percentage was computed relative to the total landing count on each day. Position landing biases were defined as the mean (across days) of landing percentages by position. We simulated the null hypothesis that observed landing counts result only from such position biases irrespective of human participants by randomly re-assigning trajectories within a given experimental day to each of the six positions, weighted according to the average position preferences (repeated 5,000 times). Statistical significance of paired rankings was evaluated based on the probability of observing an average landing percentage difference under the null hypothesis that exceeds the experimentally observed landing percentage difference.

### Weather data collection

A weather station (HOBO U30 USB) was installed outside of the cage 12m away. Information on temperature, wind speed, wind gust, relative humidity and wind direction was collected every 10 minutes for the 2020 season and 15 minutes for the 2022 season. Weather data was analyzed with MATLAB (The Mathworks Inc., R2018b).

### Seasonality of mosquito behavioral assays

All olfactory preference assays with laboratory-reared *An. gambiae* Kisumu strain in the semi-field system occurred during the wet season in Southern Zambia. Multi-choice OGTA trials with CO_2_ versus environmental air, human whole body odor versus CO_2_ and human whole body odor sourced from two humans versus environmental air occurred during the period February/March 2020. Six-choice assays with a cohort of six humans occurred during the month of April 2022.

## Supporting information

Data Sheet S1

## Acknowledgements

We thank Johns Hopkins Malaria Research Institute (JHMRI) and Bloomberg Philanthropies for generous funding to C.J.M. that supported this collaborative project with Macha Research Trust, Zambia (MRT). D.G. was supported by postdoctoral fellowships from the Human Frontier Science Program (LT000310/2019-L) and JHMRI. G.M.T received a postdoctoral fellowship from JHMRI. A.C. was supported by a graduate fellowship from the Kavli Neuroscience Discovery Institute. A.G. acknowledges funding from NIH (GM124883). We thank the JHMRI Insectary Core and Chris Kizito for *Anopheles gambiae* for lab assays, Ben Matthews for initial advice regarding infrared tracking; Marlys Book for assistance with tent modifications; Aliyah Silver for assistance with OGTA assembly; Julien Adam for drone photography; Lewis Kabinga, Pebble Moono, Saviour Simbeya, Clever Munsaka, Komana Siasikabole, Alpha Simudombe, Charlton Munsanje, Boyd Mulala and Steward Choli for assistance with the semi-field assays; the MRT entomology team for expert technical assistance and insectary support; the MRT workshop team for constructing the semi-field cage and all MRT staff for valuable logistical support.

## Author Contributions

Conceptualization, D.G., S.R.T., J.C.S. and C.J.M.; data curation, D.G., S.R.T and A.C.; formal analysis, D.G., S.R.T, A.C., A.L.G., D.M.J. and A.G.; funding acquisition: C.J.M.; investigation: D.G., S.R.T., A.L.G. and D.M.J.; methodology: D.G., S.R.T., A.C., G.M.T., A.L.G., D.M.J., C.B., A.G., and C.J.M.; project administration: L.S., C.B., J.C.S., P.T., M.M.M., E.S. and C.J.M.; visualization: D.G., S.R.T., A.C., A.G. and C.J.M.; writing – original draft, D.G., S.R.T., A.C., A.G. and C.J.M.; writing – review & editing, all authors.

---

**Supplemental Figure 1.**
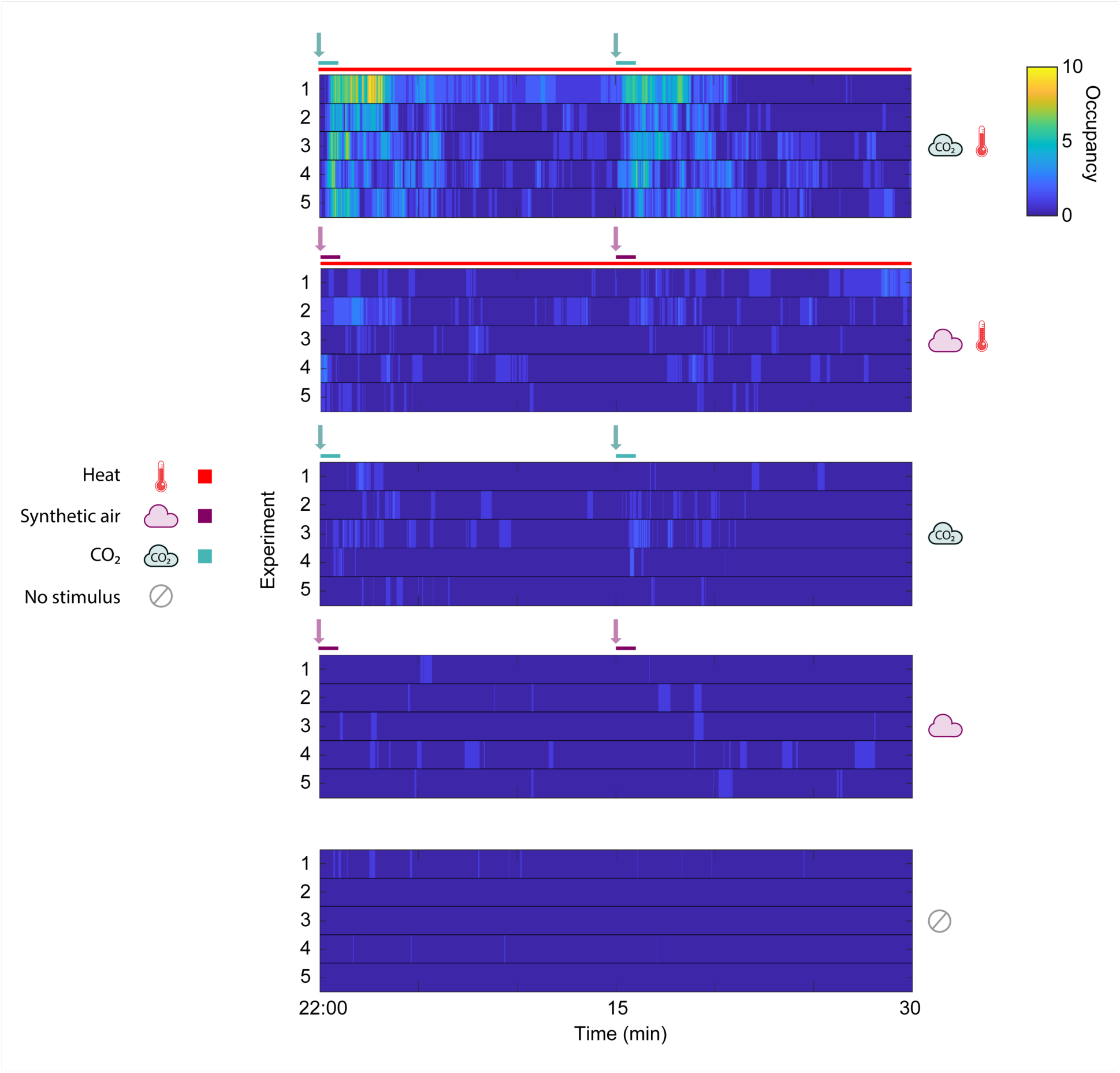
OGTA platform occupancy over time in response to combinations of multi-sensory stimuli under laboratory conditions. Occupancy represents the number of mosquitoes present on the OGTA platform in each frame (10 fps). The arrows indicate the timepoints of stimulus onset (green = CO_2_, purple = synthetic air) and the colored bars the stimulus duration. n = 5/stimulus combination. Icons represent stimulus type and combinations.

**Supplemental Figure 2.**
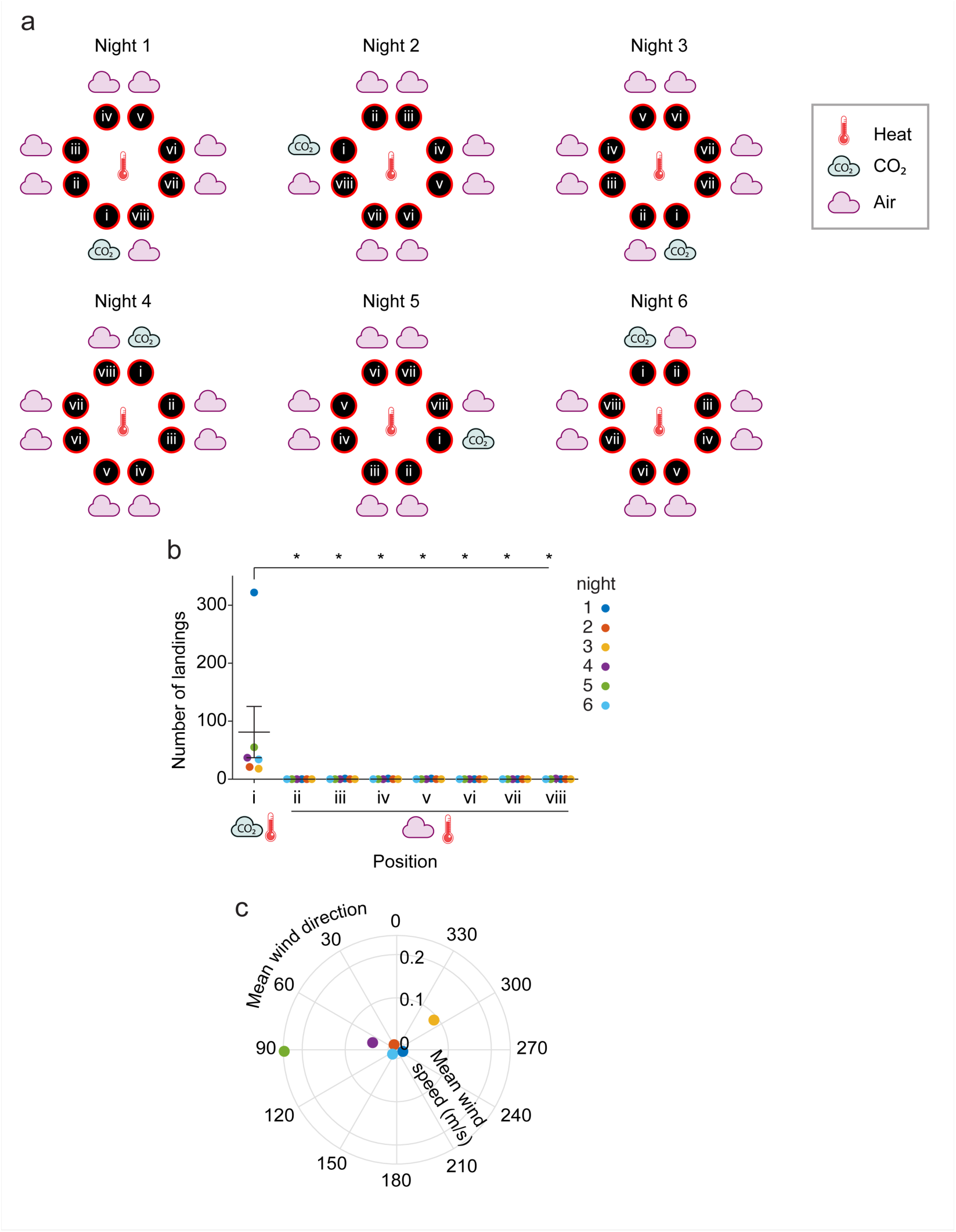
Stimulus rotation scheme, total number of mosquito landings and wind conditions for eight-choice OGTA trials with CO*_2_* versus background air under semi-field conditions. (a) Stimulus position and rotation. The CO_2_ stimulus (position i) was rotated every night. The control positions (ii-viii, air) were labeled in reference to the stimulus position in a clockwise manner. All platforms were heated to 35°C. (b) Total number of mosquito landings per night. Mean ± SEM plotted. n = 6 nights. Significant differences from CO_2_: one-sided Wilcoxon signed rank test with Benjamini-Hochberg correction (* p < 0.05). (c) Mean wind speed vs mean wind direction from 20:00 hours (time of mosquito release in cage) to 4:00 hours (end of the experiment).

**Supplemental Figure 3.**
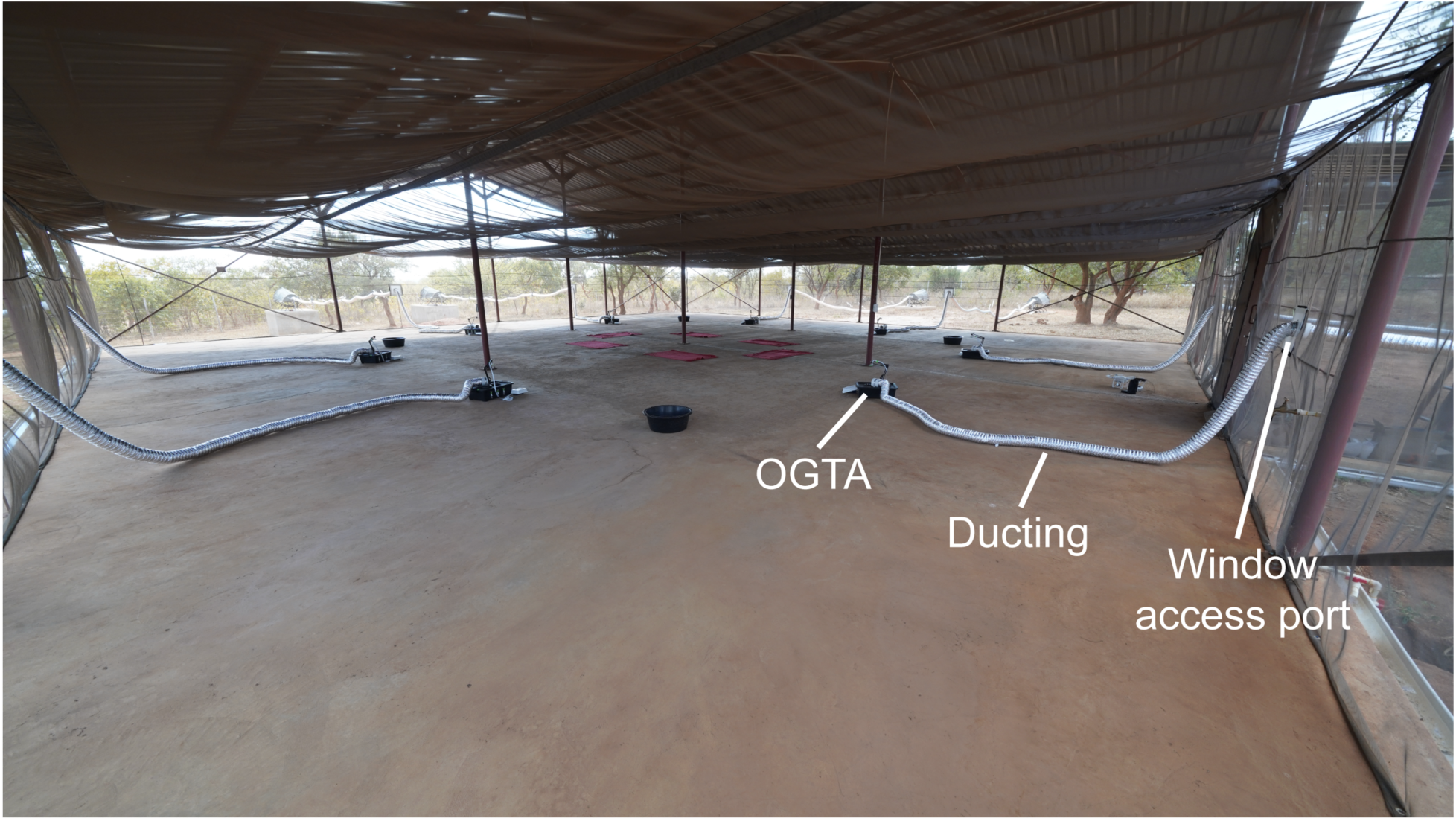
Interior view of the semi-field system with an eight-choice OGTA configuration. Eight OGTAs arranged in an octagonal array are connected to the tents outside the perimeter of the flight cage arena via ducting through widow access ports in each side of the cage.

**Supplemental Figure 4.**
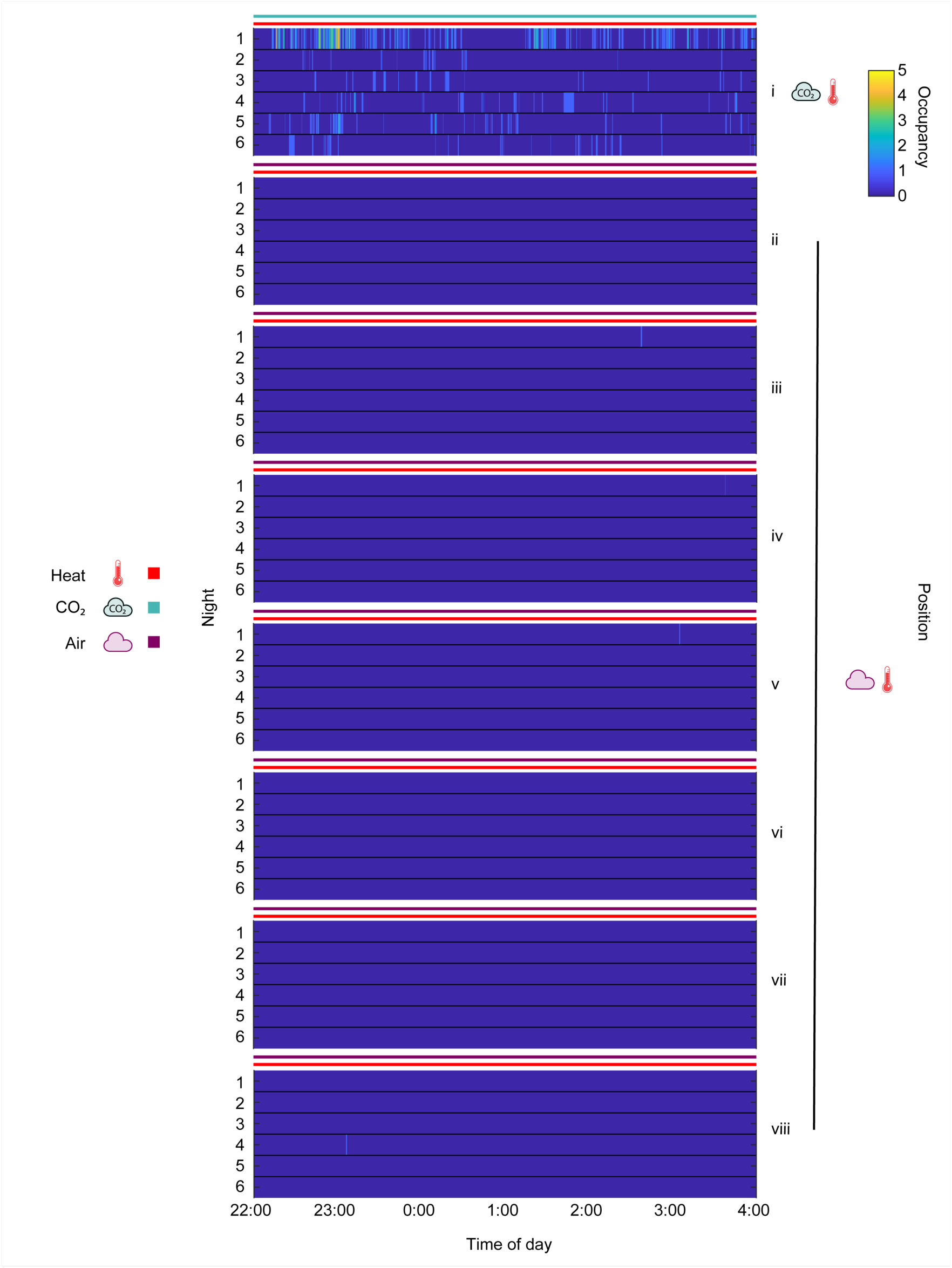
Platform occupancy over time for eight-choice OGTA trials with CO*_2_* versus background air under semi-field conditions. Occupancy represents the number of mosquitoes present on the OGTA platform in each frame (10 fps). The heat and olfactory stimuli were on for the entire duration of the experiment (colored bars). Icons represent stimulus type and combinations.

**Supplemental Figure 5.**
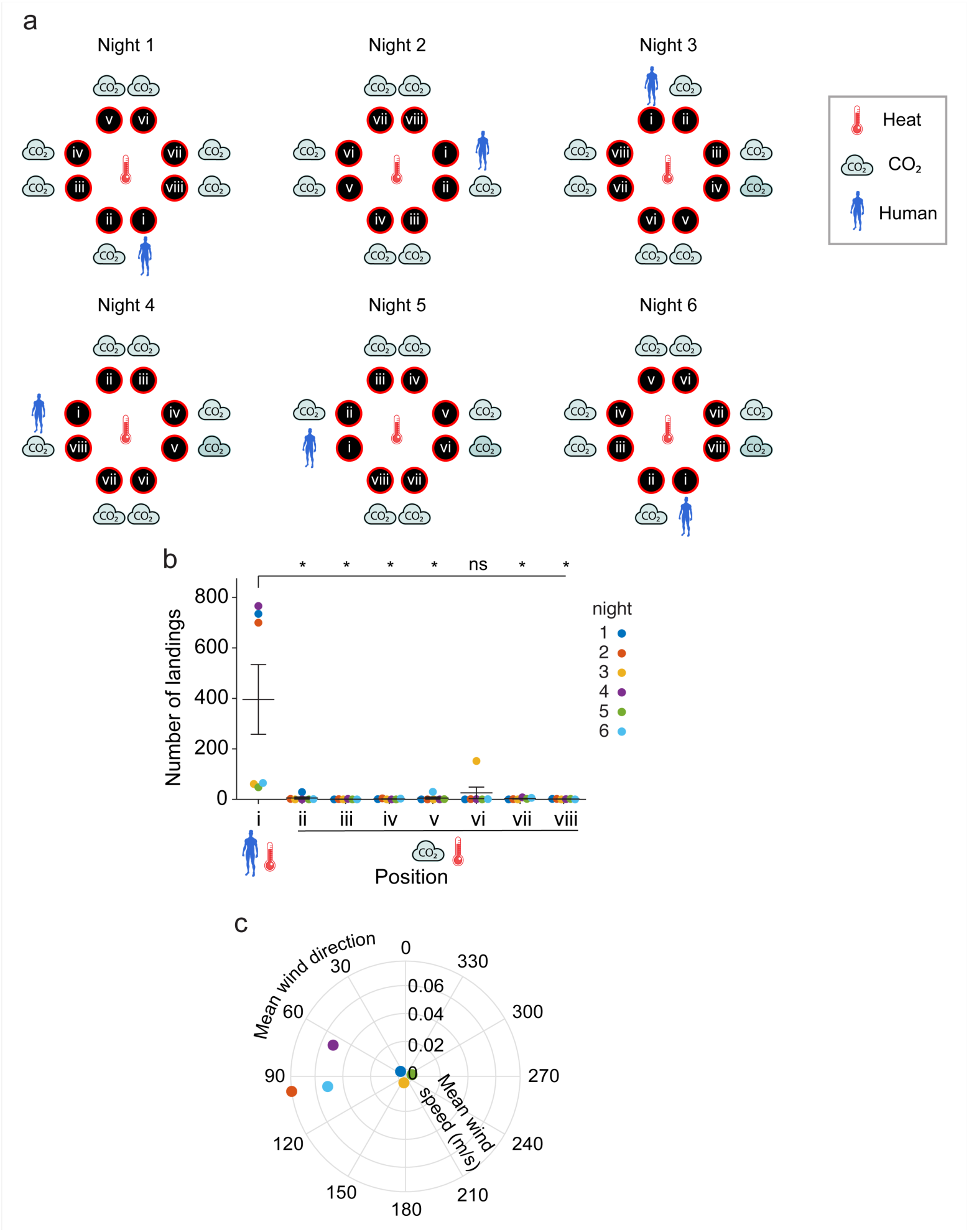
Stimulus rotation scheme, total number of mosquito landings and wind conditions for eight-choice OGTA trials with human whole body odor versus CO*_2_* under semi-field conditions. (a) Stimulus position and rotation. The human stimulus (position i) was rotated every night. The control positions (ii-viii, CO_2_) were labeled in reference to the stimulus position in a clockwise manner. All OGTA platforms were heated to 35°C. (a) (b) Total number of mosquito landings per night. Mean ± SEM plotted. n = 6 nights. Significant differences from human: one-sided Wilcoxon signed rank test with Benjamini-Hochberg correction (* p < 0.05, ns = not significant). (b) Mean wind speed vs mean wind direction from 20:00 hours (mosquito release time in the cage) to 4:00 hours (end of the experiment).

**Supplemental Figure 6.**
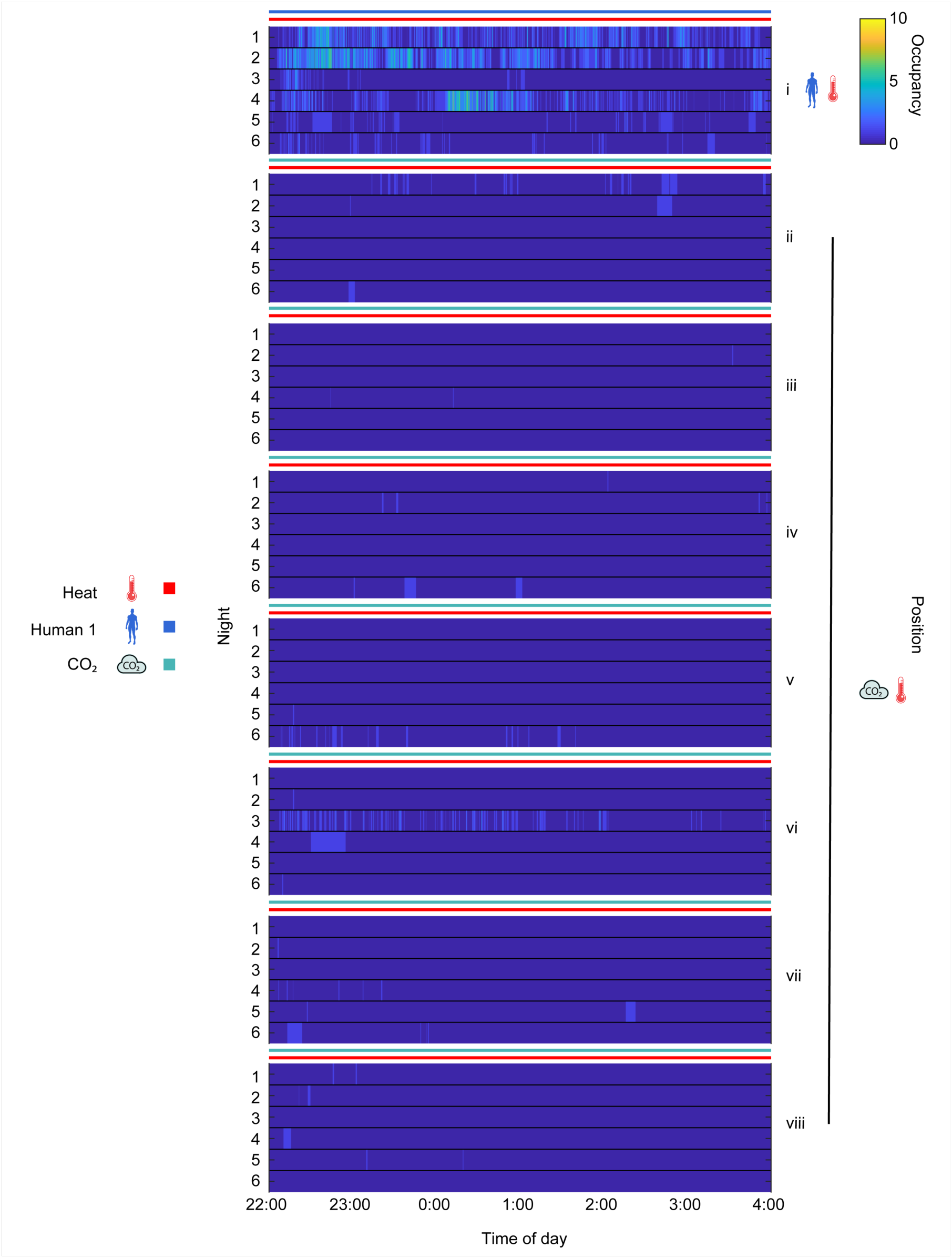
Platform occupancy over time for eight-choice OGTA trials with human body odor versus CO*_2_* under semi-field conditions. Occupancy represents the number of mosquitoes present on the OGTA platform in each frame (10 fps). The heat and olfactory stimuli were on for the entire duration of the experiment (colored bars). Icons represent stimulus type and combinations.

**Supplemental Figure 7.**
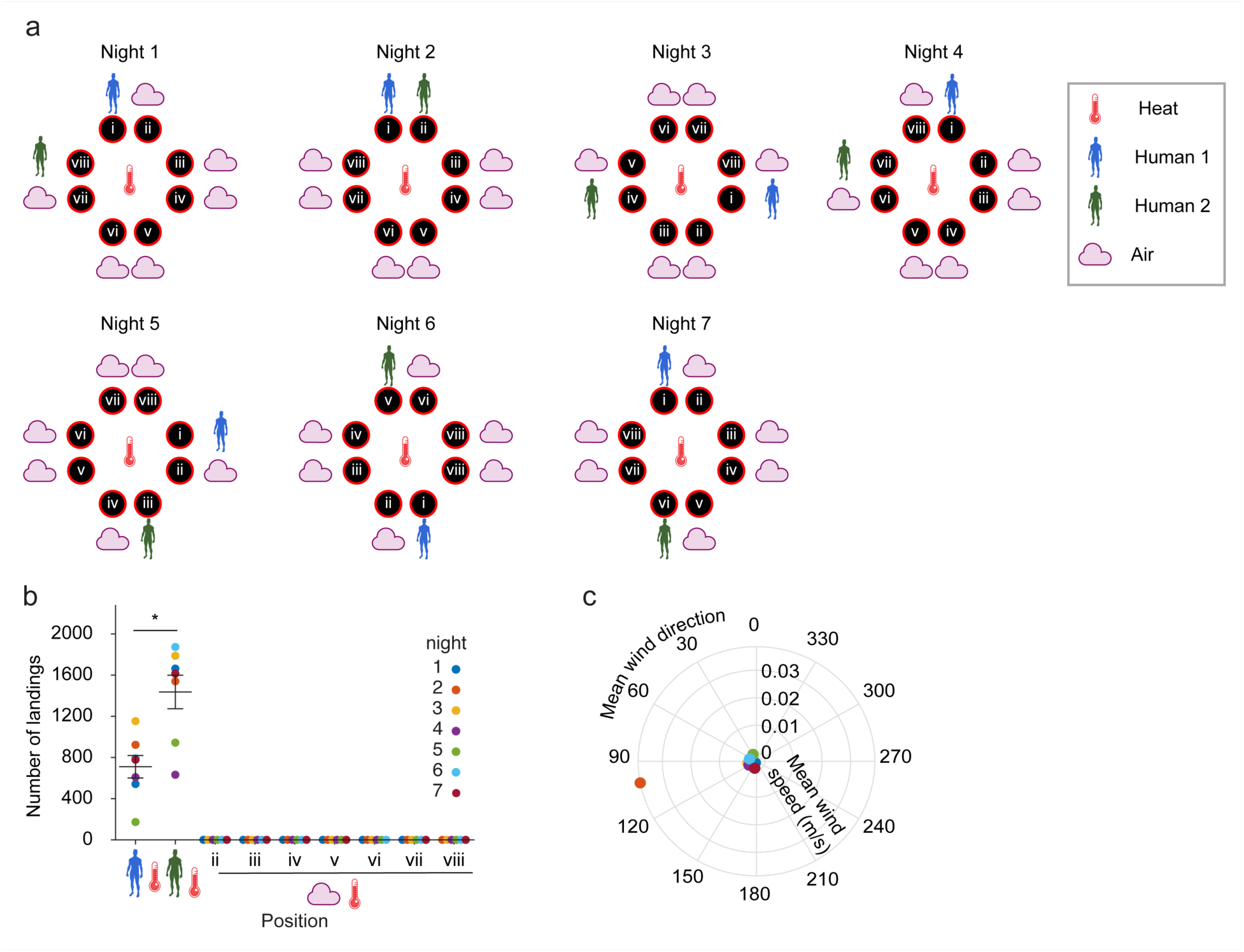
Stimulus rotation scheme, total number of mosquito landings and wind conditions for eight-choice OGTA trials with body odor from two humans versus background air under semi-field conditions. (a) Stimulus position and rotation. The position of Human 1 (blue) and the position of Human 2 (green) relative to Human 1 were rotated every night. The control positions (air) and the position of Human 2 were labeled in reference to Human 1. Human 2 could take any of the positions ii-viii. (b) Total number of mosquito landings per night. Mean ± SEM plotted. n = 7 nights (control positions have 6 data points since one of the positions was occupied by Human 2 every night). Significant differences between Human 1 and Human 2: two-sided Wilcoxon signed rank test (*p = 0.0156). (c) Mean wind speed vs mean wind direction from 20:00 hours (mosquito release time in the cage) to 4:00 hours (end of the experiment).

**Supplemental Figure 8.**
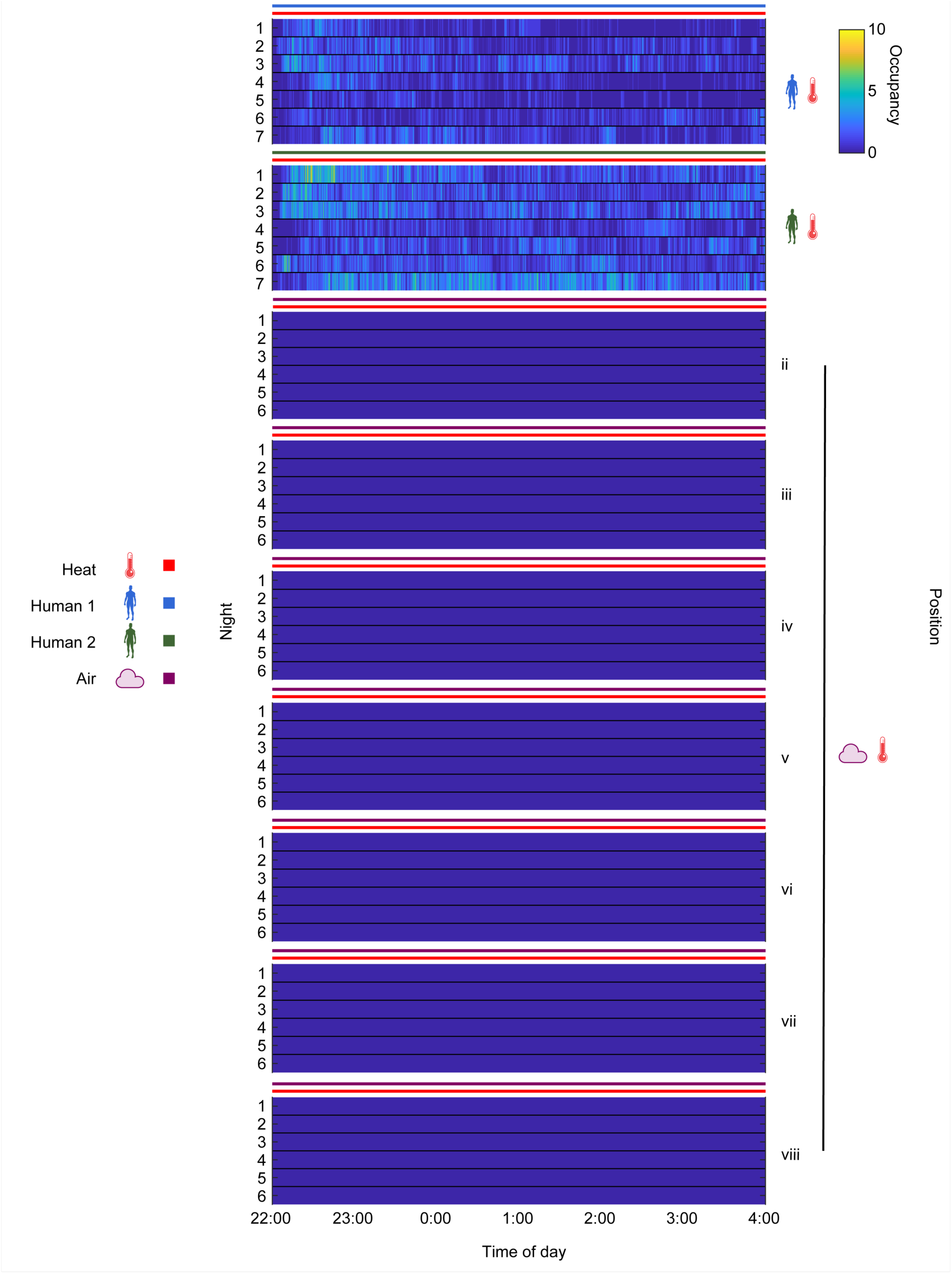
Platform occupancy over time for eight-choice OGTA trials with body odor from two humans versus background air under semi-field conditions. Occupancy represents the number of mosquitoes present on the OGTA platform in each frame (10 fps). The heat and olfactory stimuli were on for the entire duration of the experiment (colored bars). Icons represent stimulus type and combinations. Control positions have 6 nights since one of the positions was occupied by Human 2 every night.

**Supplemental Figure 9.**
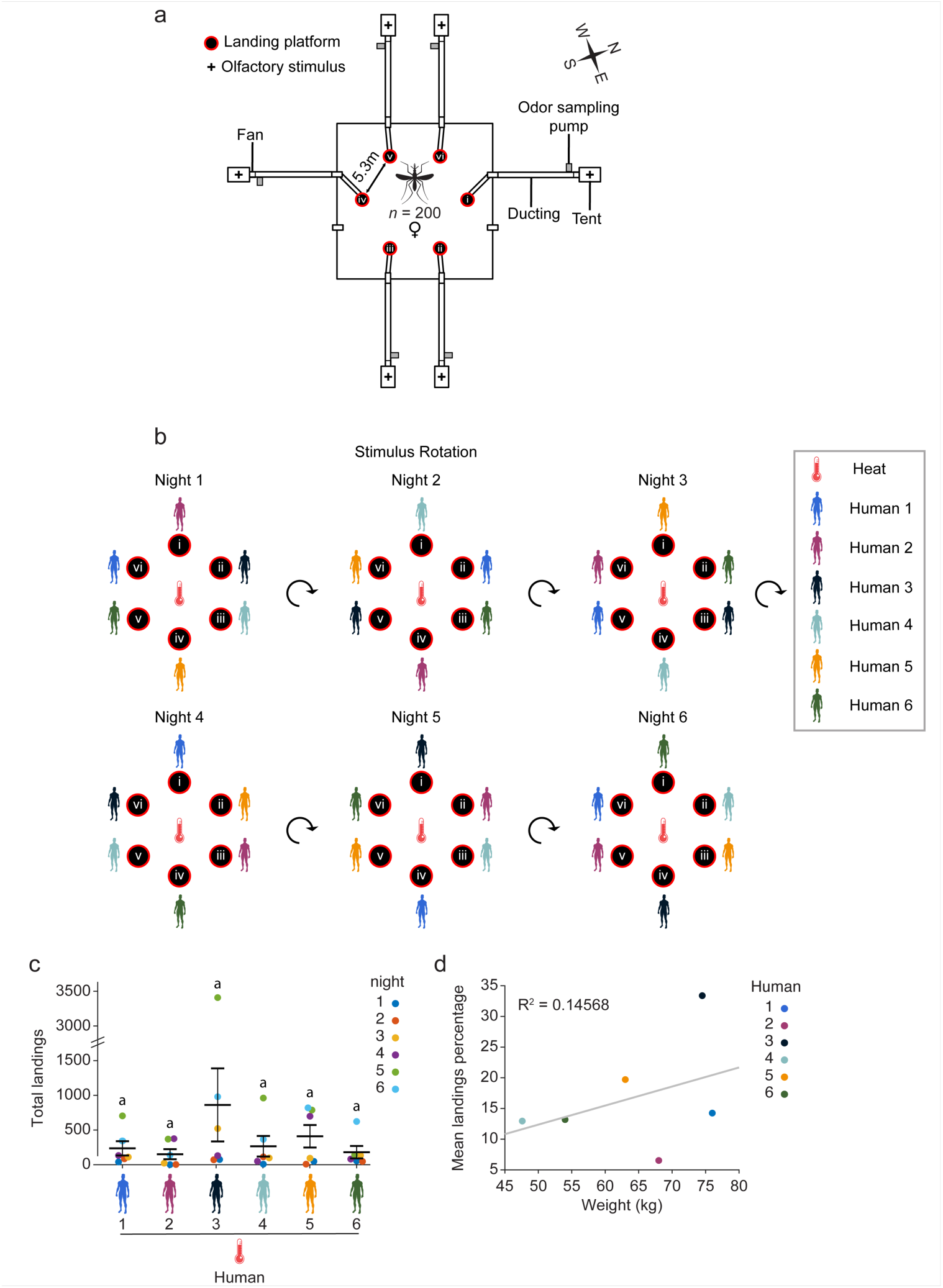
Six-choice assay configuration, stimulus rotation scheme, total number of mosquito landings and the effect of body weight on landings during trials with body odor from six humans under semi-field conditions. (a) Schematic of six-choice assay configuration. Stimulus positions were numbered relative to their position in the cage. An odor sampling pump was added to the ducting path after the fan. (b) Stimulus position and rotation for each replicate night. The positions of all humans were rotated every night. (c) Total number of mosquito landings on platforms per night per human. Mean ± SEM plotted. n = 6 nights. Significant differences between positions: Fisher’s exact permutation tests with Benjamini-Hochberg correction. (d) Participant weight versus the mean landings percentage. Coefficient of determination R^2^ = 0.14568.

**Supplemental Figure 10.**
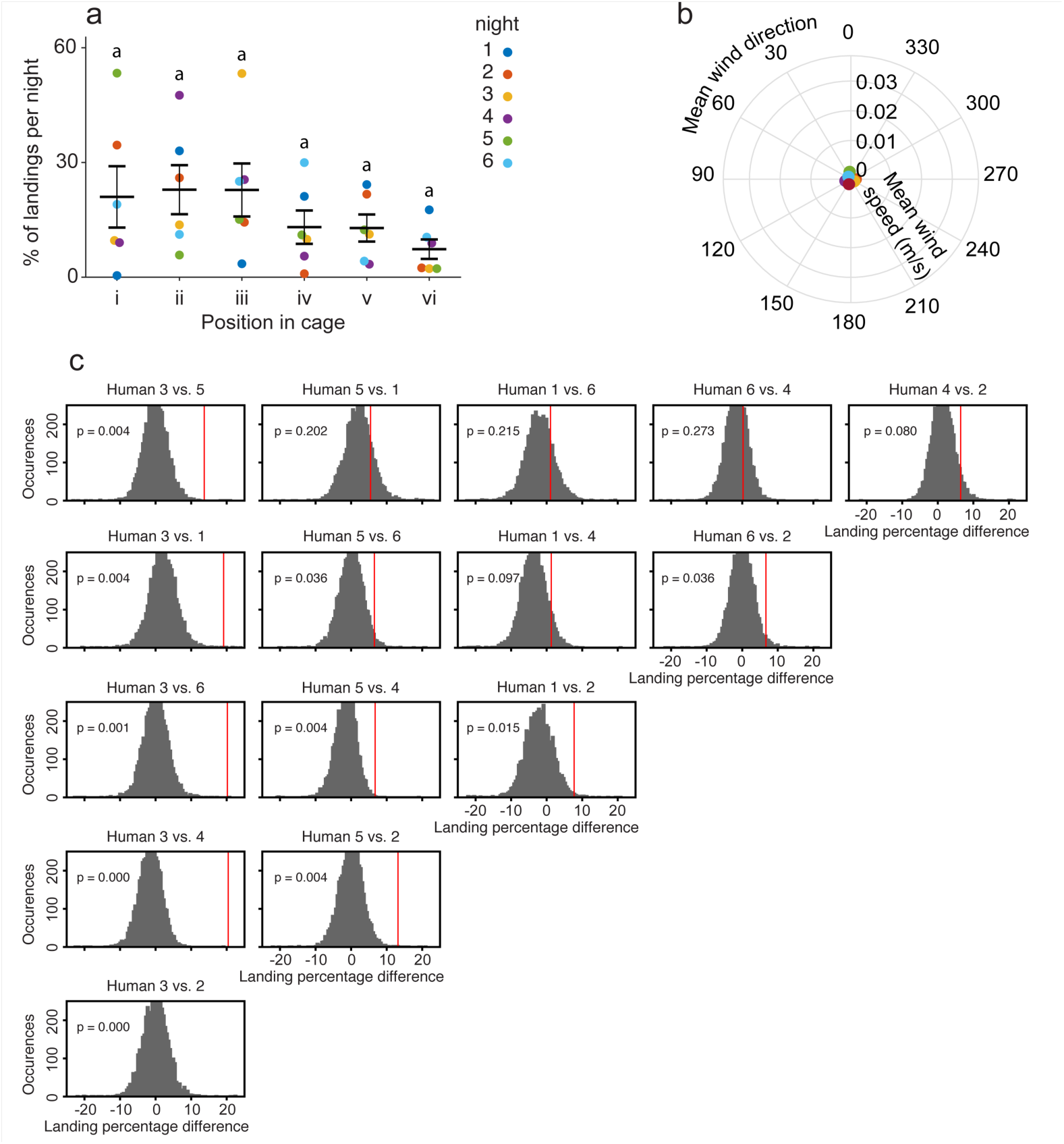
Analysis of positional effects on mosquito landings in six-choice assays. (a) Percentage landings per night per position relative to the cage in six-choice assays. Significant differences between positions: Fisher’s exact permutation tests with Benjamini-Hochberg correction. Mean ± SEM plotted. (b) Mean wind speed vs mean wind direction from 20:00 hours (mosquito release time in the cage) to 1:00 hours (end of the experiment). (c) Evaluation of the statistical significance of *An. gambiae* relative preferences for some participants over others, based on landing counts. Columns are sorted according to the mean attractiveness ranking of individuals across days. Gray bars represent the simulated probability distribution of observed landing count differences under the null hypothesis that mosquito preferences are solely driven by position preference (a) irrespective of the human participants, obtained from 5,000 simulations in which landing targets were randomly resampled based only on the average preference bias for that position. Vertical red lines indicate the observed landing percentage differences, computed within experimental days then averaged across the 6 nights. p-values for (c) indicate the probability that no behavioral preference exists for human A in favor of human B and were corrected for multiple comparisons with Benjamini-Hochberg.

**Supplemental Figure 11.**
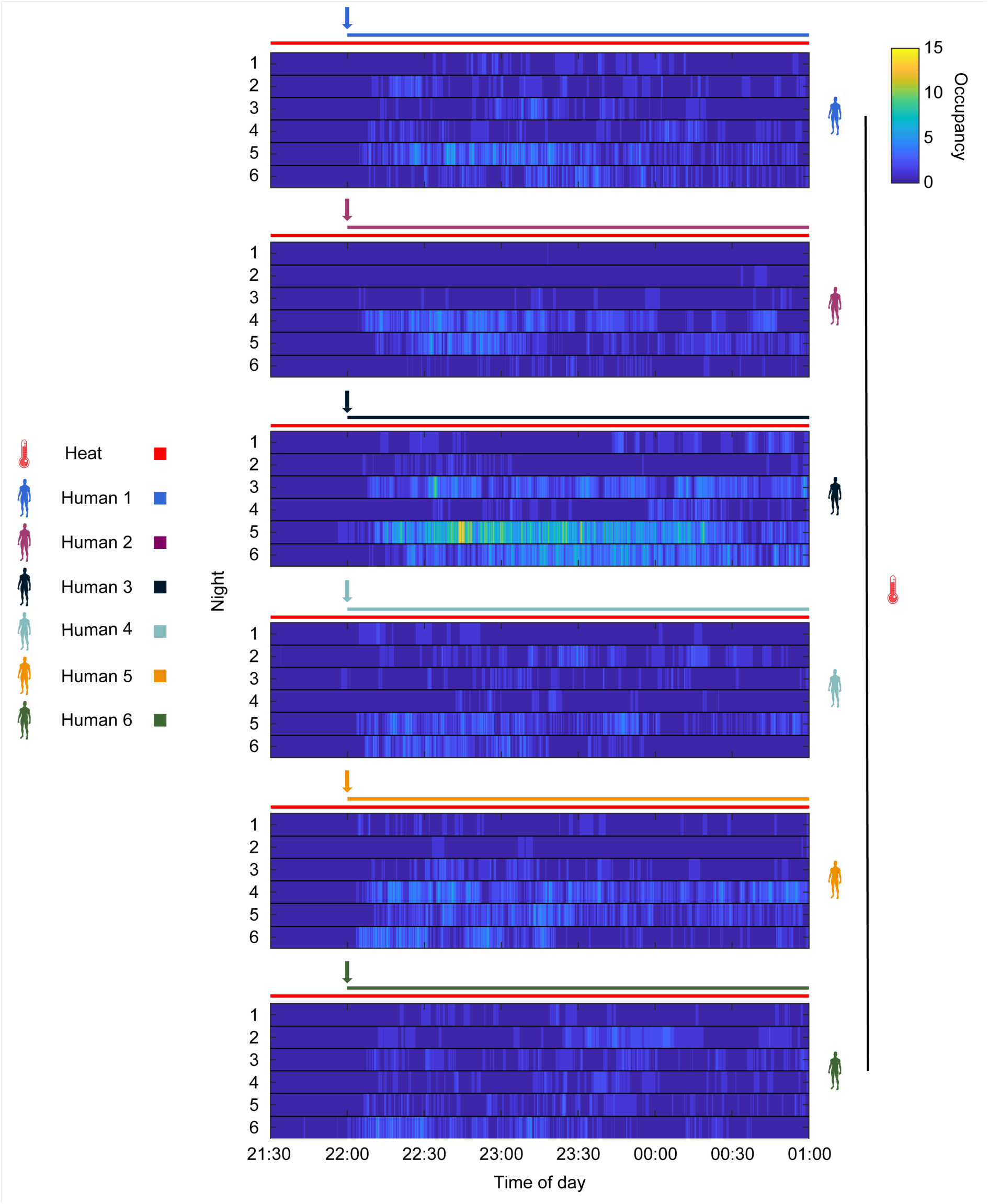
Platform occupancy over time for six-choice OGTA trials with body odor from six humans under semi-field conditions. Occupancy represents the number of mosquitoes present on the OGTA platform in each frame (10 fps). The heat stimulus was on for the entire duration of the experiment (red bars). The human stimulus was introduced at 22:00 hours (colored arrows) until the end of the experiment (colored bars). Icons represent stimulus type and combinations.

**Supplemental Figure 12.**
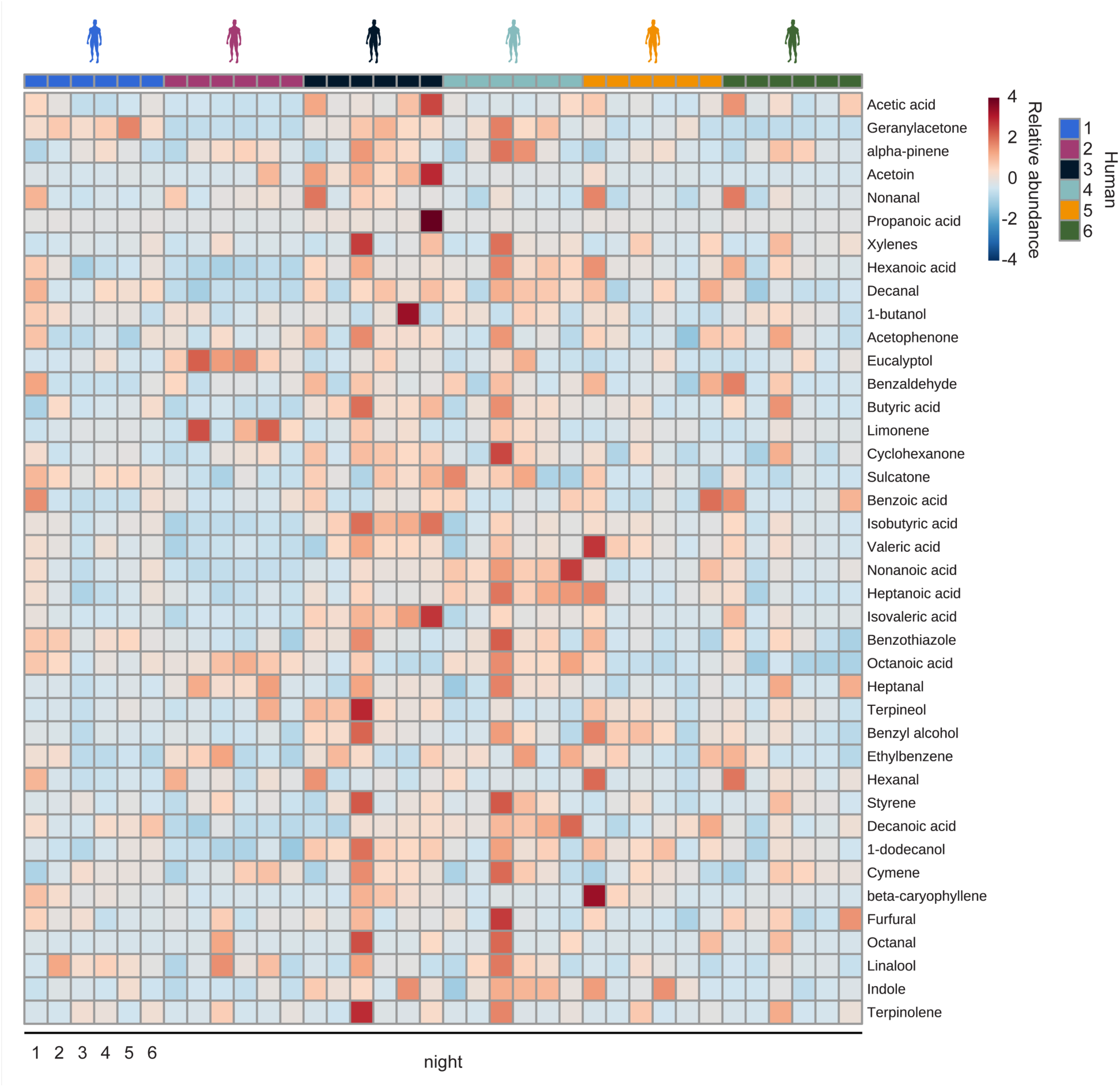
Variation in the chemical composition of human whole body odor from individuals in the six-choice assay. Heatmap of 40 identified volatile organic compounds present in human whole body odor of six human subjects, each sampled for 6 replicate nights. Scale bar represents amount of analytes detected normalized to the internal standard, with red indicating a higher concentration and blue indicating lower concentration. Heatmap constructed using MetaboAnalyst 5.0. n = 6 nights.

**Supplemental Figure 13.**
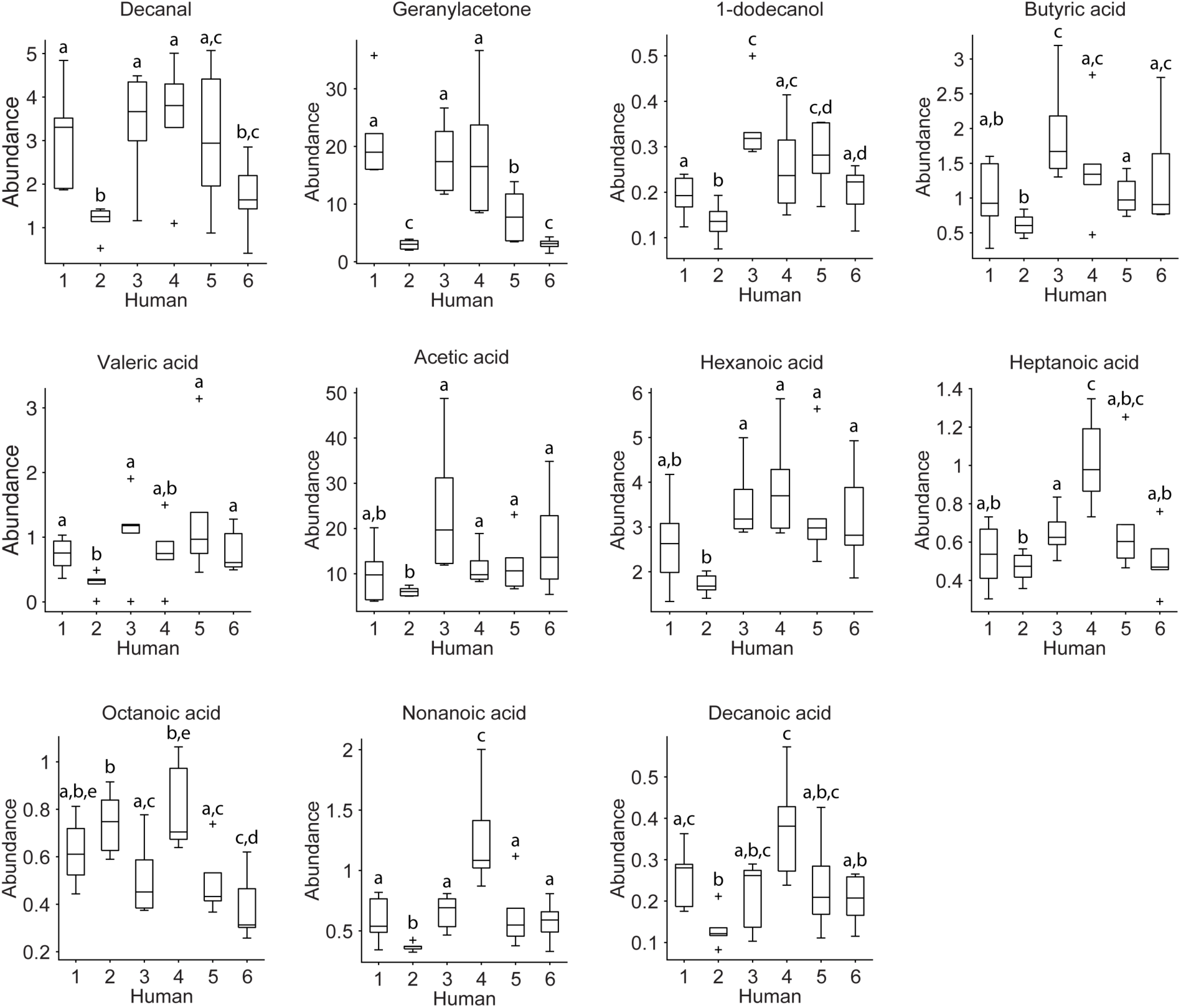
Variation in the abundance of additional human-derived volatile organic compounds from individuals in the six-choice assay. Abundances of 11/15 identified volatile organic compounds detected in whole body odor demonstrating substantial variation between participants. The line indicates the median, the box marks the lower and upper quartile, and the whiskers the 1.5 interquartile distance; outliers are indicated by black crosses. n = 6 nights. The letters indicate significant differences in compound abundance between humans: Fisher’s exact permutation tests with Benjamini-Hochberg correction.

**Supplemental Figure 14.**
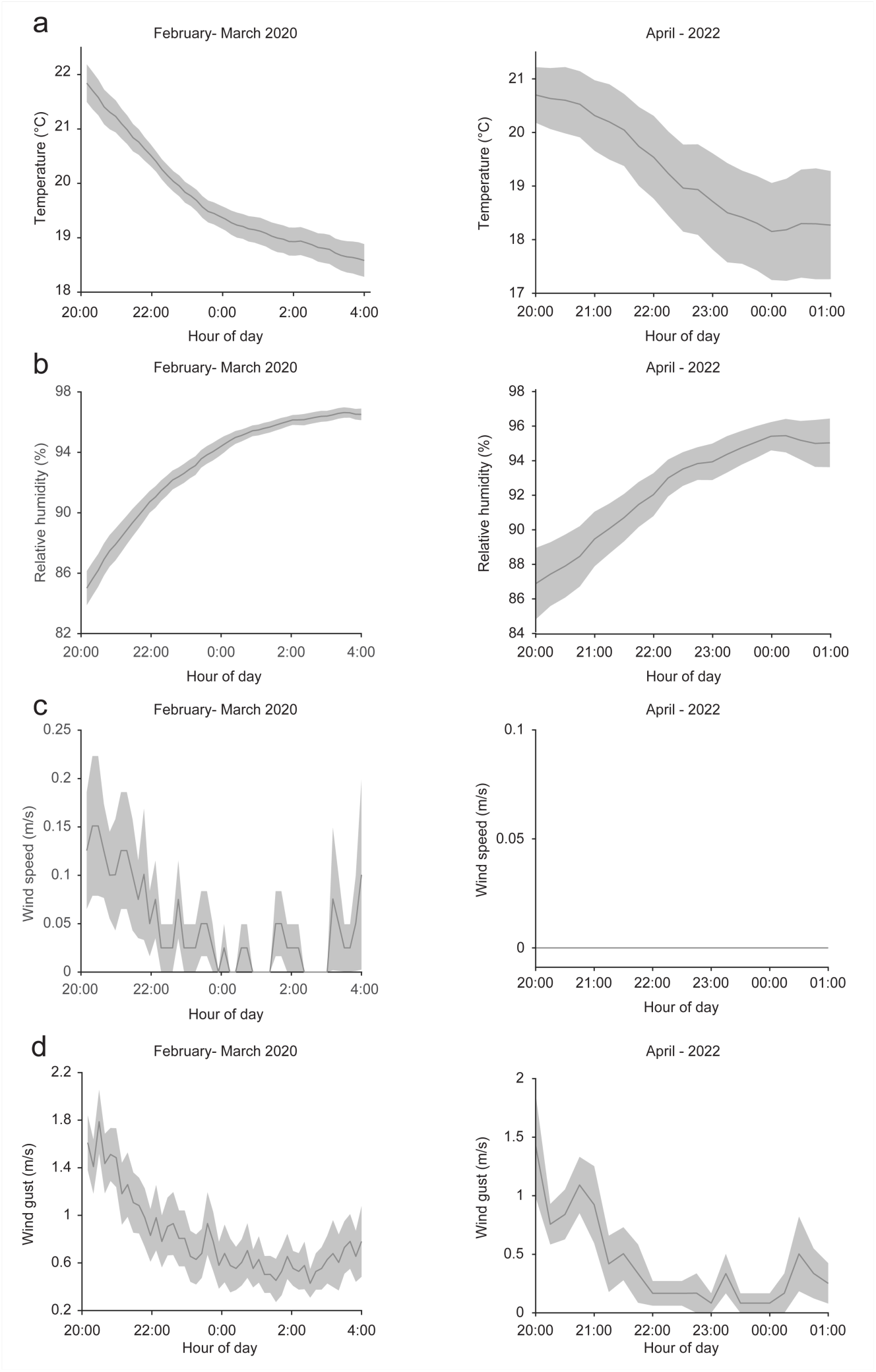
Mean weather conditions during experimental nights across test periods. Mean temperature (a), relative humidity (b), wind speed (c), and wind gust (d) for experiments carried out in 2020 (i.e. CO_2_ versus environmental air, human whole body odor versus CO_2_ and human whole body odor sourced from two humans versus environmental air) and 2022 (i.e. six-choice assays with a cohort of six humans). The data is plotted between the time of mosquito release in the cage (20:00 hours) and the end of the experiment (4:00 hours for data collected in 2020 and 1:00 hours for data collected in 2022). The data was collected by a weather station adjacent to the flight cage and averaged over the 19 total nights of experiments for 2020 and 6 for 2022. Mean ± SEM plotted.

**Supplemental Figure 15.**
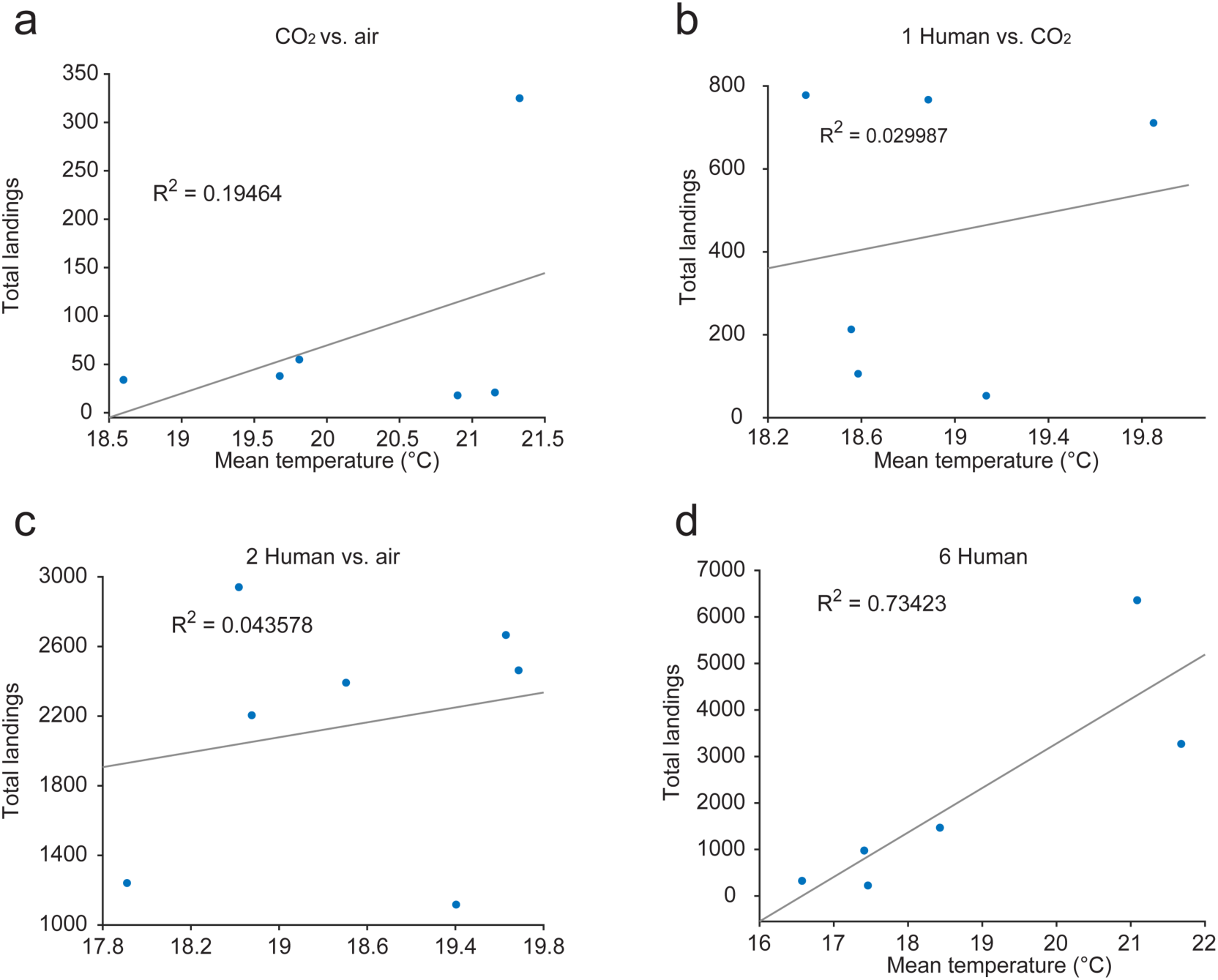
Correlation between mean temperature and total mosquito landings per night during the different semi-field experiments. (a) CO_2_ versus environmental air [CO_2_ vs. air], (b) human whole body odor versus CO_2_ [1 Human vs. CO_2_], (c) human whole body odor sourced from two humans versus environmental air [2 Human vs. air], and (d) six-choice assays with a cohort of six humans [6 Human].

## References

1. World Health Organization. World Malaria Report 2021 (2021).

2. Guelbéogo, W. M. et al. Variation in natural exposure to anopheles mosquitoes and its effects on malaria transmission. Elife 7, e32625 (2018).

3. Lindsay, S. W., Adiamah, J. H., Miller, J. E., Pleass, R. J. & Armstrong, J. R. M. Variation in attractiveness of human subjects to malaria mosquitoes (Diptera: Culicidae) in The Gambia. J. Med. Entomol. 30, 368–373 (1993).

4. Knols, B. G. J., Jong, R. de & Takken, W. Differential attractiveness of isolated humans to mosquitoes in Tanzania. Trans. R. Soc. Trop. Med. Hyg. 89, 604–606 (1995).

5. Mukabana, W. R., Takken, W., Coe, R. & Knols, B. G. J. Host-specific cues cause differential attractiveness of Kenyan men to the African malaria vector *Anopheles gambiae*. Malar. J. 1, 17 (2002).

6. Qiu, Y. T., Smallegange, R. C., Van Loon, J. J. A., Ter Braak, C. J. F. & Takken, W. Interindividual variation in the attractiveness of human odours to the malaria mosquito *Anopheles gambiae s. s*. Med. Vet. Entomol. 20, 280–287 (2006).

7. Brady, J., Costantini, C., Sagnon, N., Gibson, G. & Coluzzi, M. The role of body odours in the relative attractiveness of different men to malarial vectors in Burkina Faso. Ann. Trop. Med. Parasitol. 91, S121–S122 (1997).

8. Himeidan, Y. E., Elbashir, M. I. & Adam, I. Attractiveness of pregnant women to the malaria vector, Anopheles arabiensis, in Sudan. Ann. Trop. Med. Parasitol. 98, 631–633 (2004).

9. Smith, D. L., McKenzie, F. E., Snow, R. W. & Hay, S. I. Revisiting the basic reproductive number for malaria and its implications for malaria control. PLoS Biol. 5, 0531–0542 (2007).

10. Ansell, J., Hamilton, K. A., Pindeti, M., Walraven, G. E. L. & Lindsay, S. W. Short-range atttractiveness of pregnant women to *Anopheles gambiae* mosquitoes. Trans. R. Soc. Trop. Med. Hyg. 96, 113–116 (2002).

11. Lindsay, S. et al. Effect of pregnancy on exposure to malaria mosquitoes. Lancet 355, 1972 (2000).

12. Verhulst, N. O. et al. Cultured skin microbiota attracts malaria mosquitoes. Malar. J. 8, 302 (2009).

13. Verhulst, N. O. et al. Composition of human skin microbiota affects attractiveness to malaria mosquitoes. PLoS One 6, e28991 (2011).

14. Verhulst, N. O. et al. Differential attraction of malaria mosquitoes to volatile blends produced by human skin bacteria. PLoS One 5, e15829 (2010).

15. Shirai, Y. et al. Alcohol ingestion stimulates mosquito attraction. J. Am. Mosq. Control Assoc. 18, 91–96 (2002).

16. Lefèvre, T. et al. Beer consumption increases human attractiveness to malaria mosquitoes. PLoS One 5, e9546 (2010).

17. Paskewitz, S., Irwin, P., Konwinski, N. & Larson, S. Impact of consumption of bananas on attraction of *Anopheles stephensi* to humans. Insects 9, 129 (2018).

18. Lacroix, R., Mukabana, W. R., Gouagna, L. C. & Koella, J. C. Malaria infection increases attractiveness of humans to mosquitoes. PLoS Biol. 3, e298 (2005).

19. Robinson, A., et al. *Plasmodium*-associated changes in human odor attract mosquitoes. PNAS 115, E4209–E4218 (2018).

20. Busula, A. O. et al. Gametocytemia and attractiveness of *Plasmodium falciparum*-infected Kenyan children to *Anopheles gambiae* mosquitoes. J. Infect. Dis. 216, 291–295 (2017).

21. Fernández-Grandon, G. M., Gezan, S. A., Armour, J. A. L., Pickett, J. A. & Logan, J. G. Heritability of attractiveness to mosquitoes. PLoS One 10, e0122716 (2015).

22. Verhulst, N. O. et al. Relation between HLA genes, human skin volatiles and attractiveness of humans to malaria mosquitoes. Infect. Genet. Evol. 18, 87–93 (2013).

23. McMeniman, C. J., Corfas, R. A., Matthews, B. J., Ritchie, S. A. & Vosshall, L. B. Multimodal integration of carbon dioxide and other sensory cues drives mosquito attraction to humans. Cell 156, 1060–1071 (2014).

24. Dekker, T. & Cardé, R. T. Moment-to-moment flight manoeuvres of the female yellow fever mosquito (*Aedes aegypti* L.) in response to plumes of carbon dioxide and human skin odour. J. Exp. Biol. 214, 3480–3494 (2011).

25. Dekker, T., Steib, B., Cardé, R. T. & Geier, M. L-lactic acid: A human-signifying host cue for the anthropophilic mosquito *Anopheles gambiae*. Med. Vet. Entomol. 16, 91–98 (2002).

26. Acree Jr., F., Turner, R. B., Gouck, H. K., Beroza, M. & Smith, N. L-Lactic acid: a mosquito attractant isolated from humans. Science 161, 1346–1347 (1968).

27. Geier, M., Bosch, O. J. & Boeckh, J. Ammonia as an attractive component of host odour for the yellow fever mosquito, Aedes aegypti. Chem. Senses 24, 647–653 (1999).

28. Braks, M. A. H., Meijerink, J. & Takken, W. The response of the malaria mosquito, *Anopheles gambiae*, to two components of human sweat, ammonia and L-lactic acid, in an olfactometer. Physiol. Entomol. 26, 142–148 (2001).

29. Smallegange, R. C., Qiu, Y. T., Bukovinszkiné-Kiss, G., Van Loon, J. J. A. & Takken, W. The effect of aliphatic carboxylic acids on olfaction-based host-seeking of the malaria mosquito *Anopheles gambiae* sensu stricto. J. Chem. Ecol. 35, 933–943 (2009).

30. Kröckel, U., Rose, A., Eiras, Á. E. & Geier, M. New tools for surveillance of adult yellow fever mosquitoes: Comparison of trap catches with human landing rates in an urban environment. J. Am. Mosq. Control Assoc. 22, 229–238 (2006).

31. Mukabana, W. R. et al. A novel synthetic odorant blend for trapping of malaria and other African mosquito species. J. Chem. Ecol. 38, 235–244 (2012).

32. Okumu, F. O. et al. Development and field evaluation of a synthetic mosquito lure that is more attractive than humans. PLoS One 5, e8951 (2010).

33. Wooding, M., Naudé, Y., Rohwer, E. & Bouwer, M. Controlling mosquitoes with semiochemicals: A review. Parasites and Vectors 13, (2020).

34. Dekker, T. et al. Selection of biting sites on a human host by *Anopheles gambiae* s.s., An. arabiensis and An. quadriannulatus. Entomol. Exp. Appl. 87, 295–300 (1998).

35. Braack, L. et al. Biting behaviour of African malaria vectors: 1. Where do the main vector species bite on the human body? Parasites and Vectors 8, (2015).

36. Hinze, A., Lantz, J., Hill, S. R. & Ignell, R. Mosquito Host Seeking in 3D Using a Versatile Climate-Controlled Wind Tunnel System. Front. Behav. Neurosci. 15, (2021).

37. De Obaldia, M. E. et al. Differential mosquito attraction to humans is associated with skin-derived carboxylic acid levels. Cell 185, 4099–4116 (2022).

38. Dekker, T., Geier, M. & Cardé, R. T. Carbon dioxide instantly sensitizes female yellow fever mosquitoes to human skin odours. J. Exp. Biol. 208, 2963–2972 (2005).

39. Bernier, U. R., Kline, D. L., Allan, S. A. & Barnard, D. R. Laboratory studies of *Aedes aegypti* attraction to ketones, sulfides, and primary chloroalkanes tested alone and in combination with L-Lactic Acid. J. Am. Mosq. Control Assoc. 31, 63–70 (2015).

40. Bernier, U. R., Kline, D. L., Allan, S. A. & Barnard, D. R. Laboratory comparison of *Aedes aegypti* attraction to human odors and to synthetic human odor compounds and blends. J. Am. Mosq. Control Assoc. 23, 288–293 (2007).

41. Bello, J. E. & Cardé, R. T. Compounds from human odor induce attraction and landing in female yellow fever mosquitoes (*Aedes aegypti*). Sci. Rep. 12, (2022).

42. Khan, A. A., Maibach, H. I., Strauss, W. G. & Fenley, W. R. Screening humans for degrees of attractiveness to mosquitoes. J. Econ. Entomol. 58, 694–697 (1965).

43. Torr, S. J., Della Torre, A., Calzetta, M., Costantini, C. & Vale, G. A. Towards a fuller understanding of mosquito behaviour: use of electrocuting grids to compare the odour-orientated responses of *Anopheles arabiensis* and *An. quadriannulatus* in the field. Med. Vet. Entomol. 22, 93–108 (2008).

44. Hawkes, F. M., Dabiré, R. K., Sawadogo, S. P., Torr, S. J. & Gibson, G. Exploiting *Anopheles* responses to thermal, odour and visual stimuli to improve surveillance and control of malaria. Sci. Rep. 7, (2017).

45. Smallegange, R. C. et al. Malaria infected mosquitoes express enhanced attraction to human odor. PLoS One 8, 8–10 (2013).

46. Carnaghi, M., Belmain, S. R., Hopkins, R. J. & Hawkes, F. M. Multimodal synergisms in host stimuli drive landing response in malaria mosquitoes. Sci. Rep. 11, 7379 (2021).

47. Gillies, M. T. Age-groups and the biting cycle in *Anopheles gambiae*. A preliminary investigation. Bull. Entomol. Res. 48, 553–559 (1957).

48. Rankin-Turner, S. & McMeniman, C. J. A headspace collection chamber for whole body volatilomics. Analyst 147, 5210 (2022).

49. Lindsay, S. W. & Snow, R. W. The trouble with eaves; house entry by vectors of malaria. Trans. R. Soc. Trop. Med. Hyg. 82, 645–646 (1988).

50. Cardé, R. T. Multi-cue integration: how female mosquitoes locate a human host. Curr. Biol. 25, R793–R795 (2015).

51. Kirby, M. J. et al. Risk factors for house-entry by malaria vectors in a rural town and satellite villages in the Gambia. Malar. J. 7, 1–9 (2008).

52. Lindsay, S. W. et al. Changes in house design reduce exposure to malaria mosquitoes. Trop. Med. Int. Heal. 8, 512–517 (2003).

53. Raji, J. I. & Degennaro, M. Genetic analysis of mosquito detection of humans. Curr. Opin. Insect Sci. 20, 34–38 (2017).

54. San Alberto, D. A., et al. The olfactory gating of visual preferences to human skin and visible spectra in mosquitoes. Nat. Commun. 13, (2022).

55. Van Breugel, F., Riffell, J., Fairhall, A. & Dickinson, M. H. Mosquitoes use vision to associate odor plumes with thermal targets. Curr. Biol. 25, 2123–2129 (2015).

56. Corfas, R. A. & Vosshall, L. B. The cation channel TRPA1 tunes mosquito thermotaxis to host temperatures. Elife 4, e11750 (2015).

57. Degennaro, M. et al. *orco* mutant mosquitoes lose strong preference for humans and are not repelled by volatile DEET. Nature 498, 487–491 (2013).

58. van Loon, J. J. A. et al. Mosquito attraction: crucial role of carbon dioxide in formulation of a five-component blend of human-derived volatiles. J. Chem. Ecol. 41, 567–573 (2015).

59. Garrett-Jones, C., Boreham, P. F. L. & Pant, C. P. Feeding habits of anophelines (Diptera: Culicidae) in 1971–78, with reference to the human blood index: A review. Bull. Entomol. Res. 70, 165–185 (1980).

60. Pates, H. V., Curtis, C. F. & Takken, W. Hybridization studies to modify the host preference of *Anopheles gambiae*. Med. Vet. Entomol. 28, 68–74 (2014).

61. Mwangangi, J. M. et al. Blood-meal analysis for anopheline mosquitoes sampled along the Kenyan coast. J. Am. Mosq. Control Assoc. 19, 371–375 (2003).

62. Constantini, C., Sagnon, N. F., della Torre, A. & Coluzzi, M. Mosquito Behavioral Aspects of vector-human interactions in the *Anopheles gambiae* complex. Parasitologia 41, 209–217 (1999).

63. Dekker, T., Takken, W. & Braks, M. A. H. Innate preference for host-odor blends modulates degree of anthropophagy of *Anopheles gambiae* sensu lato (Diptera: Culicidae). J. Med. Entomol. 38, 868–871 (2001).

64. Drabińska, N. et al. A literature survey of all volatiles from healthy human breath and bodily fluids: The human volatilome. J. Breath Res. 15, 034001 (2021).

65. Takken, W. & Verhulst, N. O. Host preferences of blood-feeding mosquitoes. Annu. Rev. Entomol. 58, 433–453 (2013).

66. Bernier, U. R., Booth, M. M. & Yost, R. A. Analysis of human skin emanations by gas chromatography/mass spectrometry. 1. Thermal desorption of attractants for the yellow fever mosquito (*Aedes aegypti*) from handled glass beads. Anal. Chem. 71, 1–7 (1999).

67. Qiu, Y. T. et al. Behavioural and electrophysiological responses of the malaria mosquito *Anopheles gambiae* Giles sensu stricto (Diptera: Culicidae) to human skin emanations. Med. Vet. Entomol. 18, 429–438 (2004).

68. Roodt, A. P., Naudé, Y., Stoltz, A. & Rohwer, E. Human skin volatiles: Passive sampling and GC × GC-ToFMS analysis as a tool to investigate the skin microbiome and interactions with anthropophilic mosquito disease vectors. J. Chromatogr. B 1097–1098, 83–93 (2018).

69. Fredrich, E., Barzantny, H., Brune, I. & Tauch, A. Daily battle against body odor: Towards the activity of the axillary microbiota. Trends Microbiol. 21, 305–312 (2013).

70. Cork, A. & Park, K. C. Identification of electrophysiologically-active compounds for the malaria mosquito, *Anopheles gambiae*, in human sweat extracts. Med. Vet. Entomol. 10, 269–276 (1996).

71. Meijerink, J. & Van Loon, J. J. A. Sensitivities of antennal olfactory neurons of the malaria mosquito, *Anopheles gambiae*, to carboxylic acids. J. Insect Physiol. 45, 365–373 (1999).

72. Smallegange, R. C., Qiu, Y. T., van Loon, J. A. & Takken, W. Synergism between ammonia, lactic acid and carboxylic acids as kairomones in the host-seeking behaviour of the malaria mosquito *Anopheles gambiae* sensu stricto (Diptera: Culicidae). Chem. Senses 30, 145–152 (2005).

73. Fitzgerald, S., Duffy, E., Holland, L. & Morrin, A. Multi-strain volatile profiling of pathogenic and commensal cutaneous bacteria. Sci. Rep. 10, (2020).

74. Verhulst, N. O. et al. Improvement of a synthetic lure for *Anopheles gambiae* using compounds produced by human skin microbiota. Malar. J. 10, 28 (2011).

75. Oh, J., Byrd, A. L., Park, M., Kong, H. H. & Segre, J. A. Temporal stability of the human skin microbiome. Cell 165, 854–866 (2016).

76. Oh, J. et al. Biogeography and individuality shape function in the human skin metagenome. Nature 514, 59–64 (2014).

77. Klocke, J. A., Darlington, M. V. & Balandrin, M. F. 1,8-Cineole (Eucalyptol), a mosquito feeding and ovipositional repellent from volatile oil of *Hemizonia fitchii* (Asteraceae). J. Chem. Ecol. 13, 2131–2141 (1987).

78. Luo, D. Y., Yan, Z. T., Che, L. R., Zhu, J. J. & Chen, B. Repellency and insecticidal activity of seven Mugwort (*Artemisia argyi*) essential oils against the malaria vector *Anopheles sinensis*. Sci. Rep. 12, (2022).

79. Maia, M. F. & Moore, S. J. Plant-based insect repellents : a review of their efficacy, development and testing. Malar. J. 10, S11 (2011).

80. Henmi, A., Sugino, T., Nakamura, K., Nomura, M. & Okuhara, M. Screening of deodorizing active compounds from natural materials and deodorizing properties of cineole. J. Japan Assoc. Odor Environ. 51, 129–143 (2020).

81. European Commission, Health and Consumer Protection Directorate-General. Opinion of the Scientific Committee on Food on eucalyptol. Bruxelles-Belgium. (2002).

82. Bruderer, T. et al. On-Line Analysis of Exhaled Breath. Chem. Rev. 119, 10803–10828 (2019).

